# Enhanced surface accessibility of SARS-CoV-2 Omicron spike protein due to an altered glycosylation profile

**DOI:** 10.1101/2023.11.22.568381

**Authors:** Dongxia Wang, Zijian Zhang, Jakub Baudys, Christopher Haynes, Sarah H. Osman, Bin Zhou, John R. Barr, James C. Gumbart

## Abstract

SARS-CoV-2 spike (S) proteins undergo extensive glycosylation, aiding proper folding, enhancing stability, and evading host immune surveillance. In this study, we used mass spectrometric analysis to elucidate the N-glycosylation characteristics and disulfide bonding of recombinant spike proteins derived from the SARS-CoV-2 Omicron variant (B.1.1.529) in comparison with the D614G spike variant. Furthermore, we conducted microsecond-long molecular dynamics simulations on spike proteins to resolve how the different N-glycans impact spike conformational sampling in the two variants. Our findings reveal that the Omicron spike protein maintains an overall resemblance to the D614G spike variant in terms of site-specific glycan processing and disulfide bond formation. Nonetheless, alterations in glycans were observed at certain N-glycosylation sites. These changes, in synergy with mutations within the Omicron spike protein, result in increased surface accessibility of the macromolecule, including ectodomain, receptor-binding domain, and N-terminal domain. These insights contribute to our understanding of the interplay between structure and function, thereby advancing effective vaccination and therapeutic strategies.

**Teaser:** **Through mass spectrometry and molecular dynamics simulations, SARS-CoV-2 Omicron spike is found to be less covered by glycans when compared to the D614G spike variant.**

## Introduction

The COVID-19 pandemic, caused by the severe acute respiratory syndrome coronavirus 2 (SARS-CoV-2), continues to pose a significant global health challenge (*1-3*). Since its appearance, the virus has been evolving through multiple genetic changes, leading to the emergence of variants with diverse characteristics (*4-6*). Omicron is the most recently circulating variant and its subvariants are still the dominant strains spreading worldwide during 2023 (*7, 8*). The Omicron lineages have attracted substantial attention due to their unusually high number of mutations (Fig. 1) in the spike protein, which has resulted in increased transmissibility, milder symptoms, and low severity in infected patients (*9, 10*).

**Fig. 1.**
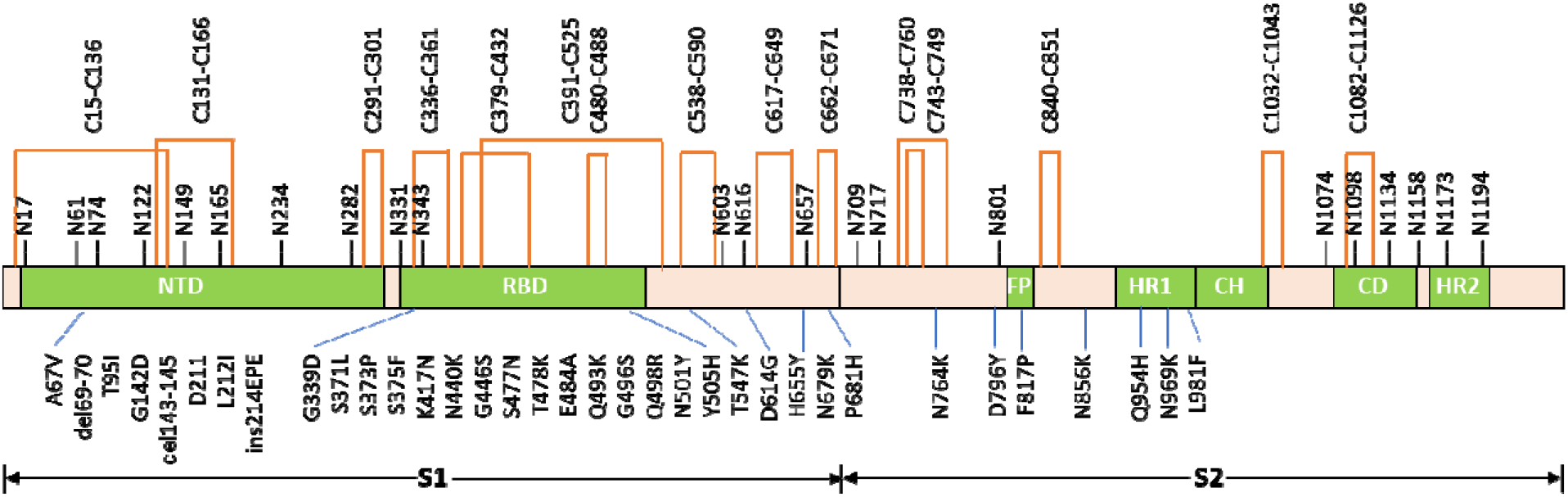
Schematic structure of the SARS-CoV-2 Omicron (B.1.1.529) spike protein, including amino acid substitution (below), N-glycosylation sites (above), and disulfide bonds (top). Domains include: the N-terminal domain (NTD); receptor binding domain (RBD); fusion peptide (FP); heptad repeat1 (HR1): central helix (CH) region; connector domain (CD); and heptad repeat 2 (HR2).

The spike protein (S) of SARS-CoV-2 virus plays a pivotal role in viral entry and infection (*11, 12*). It has been characterized that this membrane anchored S protein forms a homotrimer structure and each of the protomers contains a S1 and S2 subunits responsible for different functions (*13, 14*). The S1 subunit comprises the N-terminal domain (NTD) and receptor binding domain (RBD), a key component mediating the attachment of the virus to the host cell receptor angiotensin-converting enzyme 2 (ACE2), while the S2 subunit, containing the fusion peptide and other domains, is responsible for viral membrane fusion (*12, 15, 16*). The spike glycoprotein is the target for neutralizing antibodies and therefore is the primary target for vaccine development and therapeutic interventions (*12, 17*).

N-glycosylation, the attachment of carbohydrate moieties to asparagine residues, is a vital post-translational modification that influences protein folding, stability, and immune recognition (*18*). The S protein of SARS-CoV-2 virus is heavily glycosylated with 22 potential N-glycosylation sites (sequons, Fig. 1) on each S protomer. These glycans can shield antigenic sites from immune surveillance, affecting viral neutralization and immune evasion strategies (*19-22*). Previous studies have also demonstrated that the glycans play other roles beyond shielding the S protein from host immune recognition (*23, 24*). Since the outbreak of the COVID-19 pandemic, the glycosylation profile of the SARS-CoV-2 S glycoproteins on the initial strain and evolving variants has been intensively studied, particularly through the use of advanced liquid chromatography coupled to tandem mass spectrometry (LC-MS) techniques (*20, 23, 25-41*) with recombinantly expressed proteins of ectodomain or subunit constructs and even viral derived S protein (*14*). The analysis of glycopeptides derived from S proteins allows the determination of N-glycan profiles for all 22 conserved sequons as well as two novel sequons in Gamma spike protein.

Disulfide bonds are another important modification critical for maintaining the structural integrity of proteins. These covalent bonds between cysteine residues contribute to protein folding, stability, and overall conformation (*42*). The primary sequence of the ectodomain of the SARS-CoV-2 S protein has 30 cysteine residues (Fig. 1). Structural and mass spectrometry analysis have revealed that these cysteines form 15 disulfide bonds (*12, 13, 40*). Disulfide bonds in S proteins play an important role in their structure and function. Studies have shown that thiol-based drugs can impair the binding of S protein to ACE2 (*43-45*), and engineered disulfide bonds can trap S protein in an RBD “down” conformation (*46, 47*). The disulfide bond between C840 and C851 in the fusion peptide of S protein also can facilitate the binding between this peptide and cell membrane (*48*). A recent study has revealed that mutations in the RBD domain of the Omicron S protein affect the stability of two disulfide bonds, elevating the vulnerability of this S variant to reduction (*49*).

Our aim in the present study was to better understand the structural characteristics and changes of the SARS-CoV-2 Omicron S protein (S-Omicron) relative to the S protein of the D614G variant (S-D614G). To do that, we used LC-MS to analyze the N-glycosylation profile and disulfide bonds for each, revealing distinct distributions of glycans at some sequons. Molecular dynamics simulations of models of S-Omicron and S-D614G trimers based on the LC-MS results further elucidated changes in protein conformational sampling as a result. In summary, we provide insights into the potential consequences of structural changes in variants on viral structure, immune recognition, and therapeutic strategies.

## Results

### Characterization of N-Glycosylation on S proteins

Multiple enzyme digestion is a method commonly used to characterize post-translational modifications by the “bottom-up” MS/proteomics approach. Because of nonspecific cleavage and production of undetected short or long peptides, sequence coverage of a protein characterized by this technique is usually limited by digestion with a single protease, such as trypsin, thus hindering identification and quantification of the peptides bearing target post-translational modifications. Microheterogenity (multiple glycans on one site) of glycan distribution on glycoproteins adds to the difficulty of site-specific glycosylation analysis. Therefore, almost all researchers, including us, have used two or more proteases to digest proteins to generate peptides with a single glycosylation site and uniform sequences for characterizing glycosylation on SARS-CoV-2 spike proteins (*27-29, 31-33, 35, 37-41, 50*). During the analysis of N-glycosylation of some SARS-CoV-2 spike variants of concern, we became aware that different enzyme sets could generate results with significant variation for a single sequon. We therefore developed a set of criteria to select the best dataset for individual glycosylation sites (*37*). Similar approaches were applied in this study. Five different enzyme combinations, including Lys-C/Lys-C, Lys-C/trypsin, Lys-C/chymotrypsin, Lys-C/a-litic protease, and Asp-N/Chymotrypsin, were able to provide optimal results of the N-glycosylation profiles at all 22 sites (Table S1). As expected, consistent glycan distributions were obtained from the glycopeptides derived from different enzyme digests for many sequons. However, some digestion pairs generated inconsistent results for the same site. For instance, three digestions with Lys-C/Lys-C, Lys-C/trypsin, and Lys-C/chymotrypsin yielded similar results in terms of the number of N-glycans and the relative abundance of the glycans of different processed level for N282, but the other set of data obtained from Lys-C/alphalitic protease showed significant differences (Table S1). Our data also showed that the optimal protease combination for a specific sequon on S-Omicron was not necessarily the best one for other SARS-CoV-2 variants of concern spikes, presumably because the different amino acid substitutions among these variants alters the accessibility of some enzymatic cleavage sites altering the efficiency of the enzymes. These results underline the importance of applying multiple protease digests and obtaining consistent data from at least two enzyme combinations whenever possible for confirmation.

Fig. 2 shows the distribution of different types of glycans and their contents of fucose and sialic acid groups on all individual sequons of the Omicron spike trimer and the control sample, S-D614G. The majority of the 22 sequons in both S-Omicron and S-D614G were occupied predominantly by either underprocessed oligomannose (2 HexNAc and greater than 4 Hex groups) or fully processed complex (more than 3 HexNAc and various Hex) glycans. Whereas complex glycans occupied approximately half or more of the sites N17, N74, N149, N331, N343, N657, N1158, N1173 (S-Omicron) and N1194, the oligomannose glycans dominated the sites of N61, N122, N165 (in S-Omicron), N234, N603, N616, N709, N717, N801, N1074, and N1098 (S-Omicron). A few sites, including N1098 and N165, were detected to have approximately half of hybrid (3 HexNAc) glycans. Approximately half of the N1173 and N1194 sites were not occupied, and very low level of paucimannose (1-2 HexNAc and less than 4 Hex) was detected at some sequons.

**Fig. 2.**
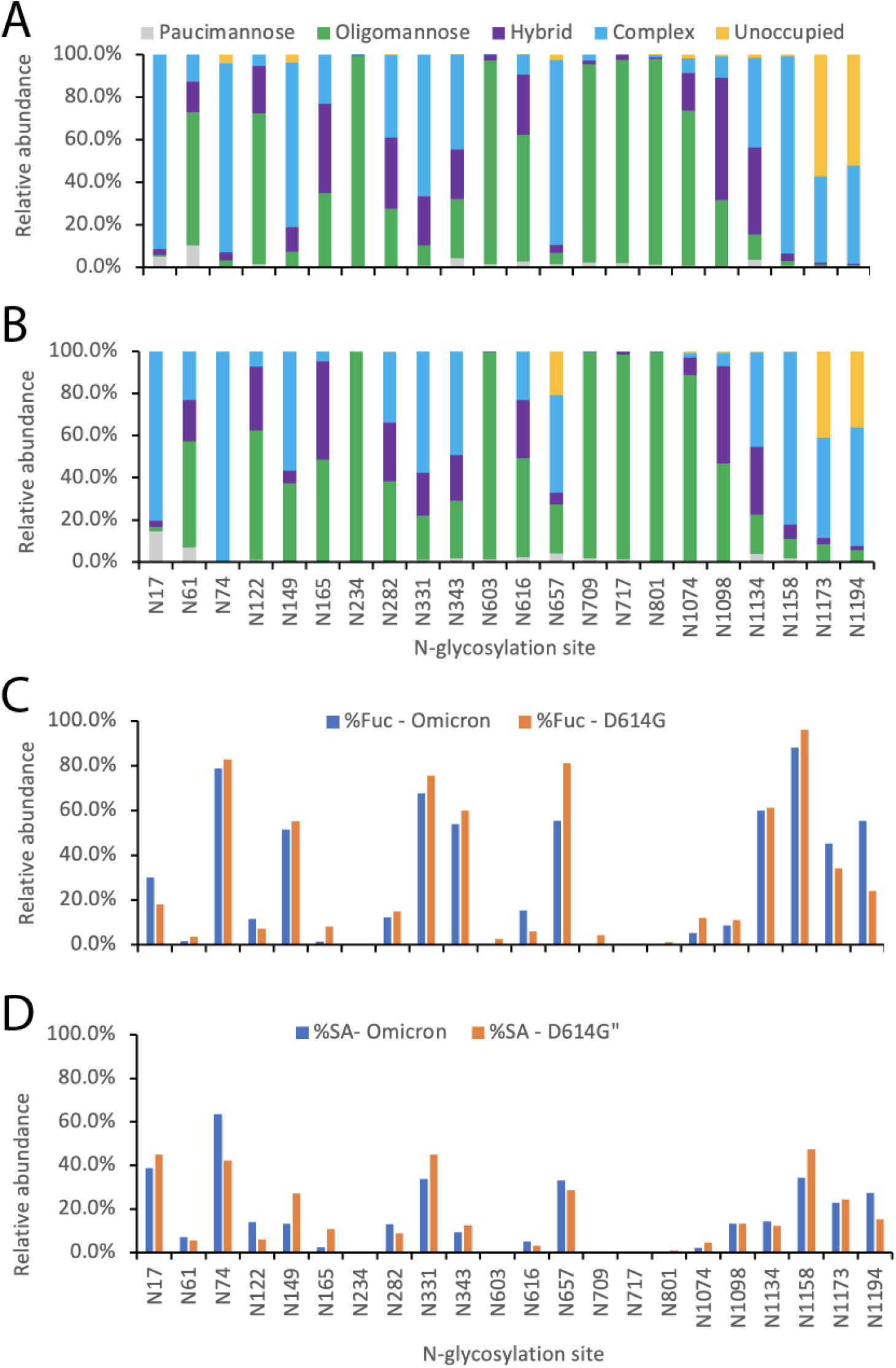
N-glycosylation profiles of S-D614G (A) and S-Omicron (B), and the relative abundance of fucosylated (Fuc) (C) and sialylated (SA) glycans (D) in these two proteins.

Inconsistent or conflicting results regarding the N-glycosylation profile of the ectodomain SARS-CoV-2 S protein have been reported from various studies (*27*). This might be caused partially by the location of the sequons and subtle changes on the structure of the trimeric S protein as well as the conditions that made the recombinant proteins. For example, in terms of oligomannose content, this report and all previous studies have determined that N234 is occupied predominantly by this type of glycan. This is true no matter the differences in protein source, type of variants, MS techniques, and data processes used in these studies, presumably due to the relatively buried location of this residue (*13*). Using human-cell-expressed ectodomain S protein, many laboratories (including ours in this study) have determined that some other sequons, such as N61, N603, N709, N717, N801, and N1074, are heavily occupied by oligomannose-type glycans (*14, 25, 26, 30, 31, 39, 41*). The fact that these residues are in the area between the head and the stalk of an S trimer structure might suggest that steric effects exist in these areas, preventing the access of glycosylation enzymes. On the other hand, two recent reports have revealed that only N61 and N234 are almost fully modified by oligomannose glycans on the S variants investigated (*33, 37*). This might be because all the proteins in these two studies were obtained from the same manufacturer. The constructs or protein preparation conditions of these samples also might be different from other sources, leading to subtle structure alterations in the middle area between the head and stalk of the S trimer.

Previous reports from various laboratories have consistently showed a high degree of similarity in the N-glycosylation profile of human embryonic kidney (HEK)-cell-expressed recombinant ectodomains of the spike proteins from different variants of concern, including Wuhan-hu-1, D614G, Alpha, Beta, Gamma, Delta, Omicron, and spike protein derived from the virus itself (*14, 26, 31, 33, 37*). This similarity was also observed in our study when comparing the glycan patterns of S-Omicron and S-D614G (Fig. 2A and 2B). Additionally, we found comparable relative abundance of fucosylated and sialylated glycans (Fig. 2C and 2D), along with the similar distribution of the most abundant glycans (Fig. 3), further indicating that the virus might have almost fully optimized the potential of N-glycosylation to evade immune responses (*31*).

**Fig. 3.**
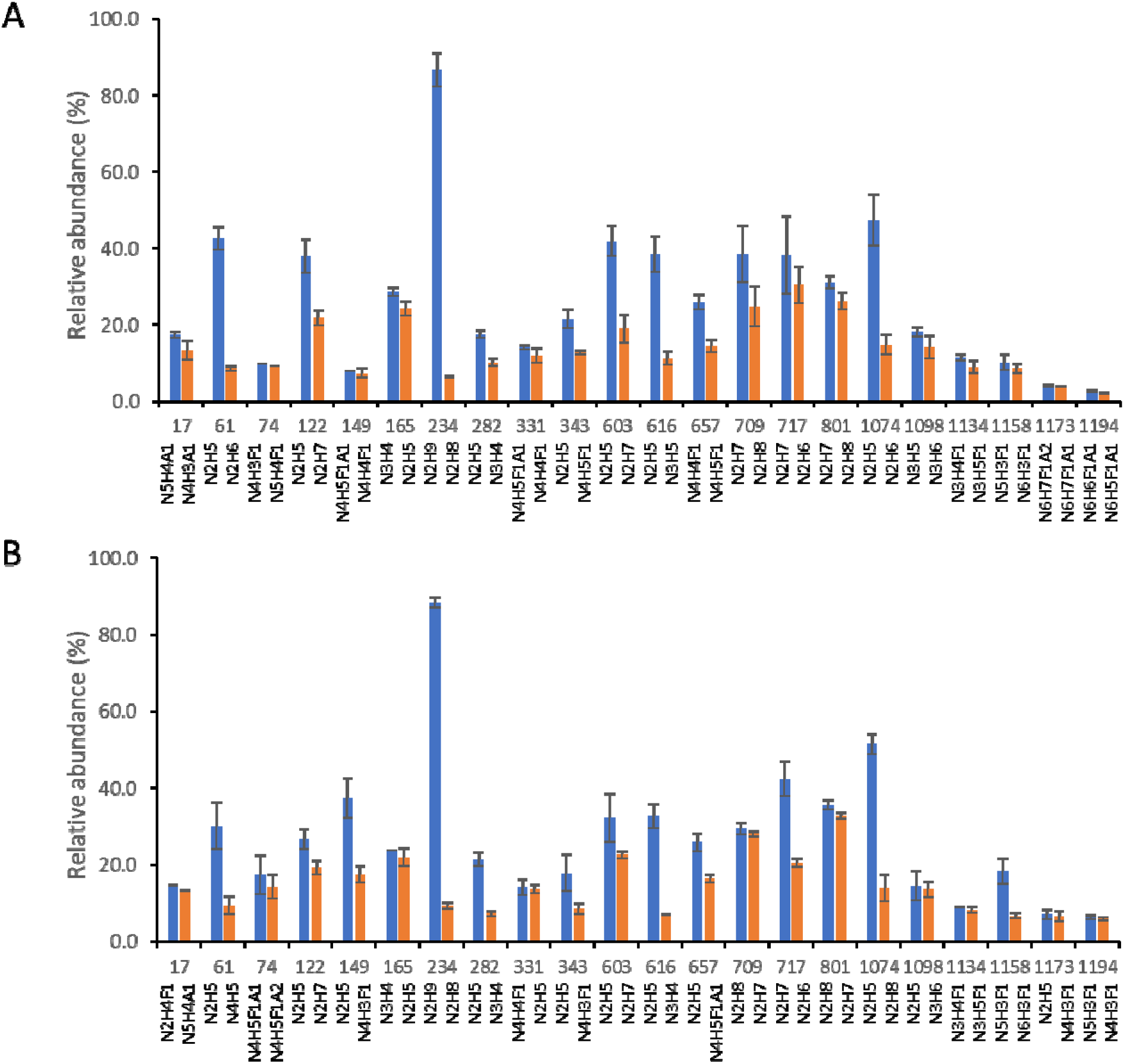
Relative abundance (%) of the most abundant (blue bar) and next most abundant (orange bar) glycans at individual N-glycosylation sites of the spike proteins of SARS-CoV-2 D614G (A) and Omicron (B) variants. N, H, F, and A represent N-acetyl hexosamine (HexNac), hexose (Hex), fucose (Fuc), and N-acetylneuraminic acid (NeuAc) groups, respectively, in glycan compositions.

Despite the overall similarity of the N-glycosylation profiles between S-Omicron and S-D614G, significant differences were observed for some sequons. For example, the content of the complex glycans on N149 decreased from 77% on S-D614G to 57% in S-Omicron, whereas the oligomannose increased from about 7% to 37%, and the major oligomannose glycan of HexNAc2Hex5 displayed approximately 40% occupancy at this site of S-Omicron (Fig. 3B). This might be attributed to the variant-specific mutations on S-Omicron near N149, including G142D and del143-145 (Fig. 1), which create a steric microenvironment preventing the access of glycan processing enzymes to this residue. A similar trend was observed at the N657 site, where the complex glycans decreased from 87% in S-D614G to 46% in S-Omicron and the oligomannose increased by approximately 4-fold. In addition, the two most abundant glycan groups at N657 were different in these two proteins (Fig. 3). A substitution in S-Omicron on a nearby residue, H655Y, might contribute to this shift. On the other hand, other N-glycosylation sites near variant-specific mutations on S-Omicron did not cause significant change in the processing state of modified N-glycans. For instance, the profile of the glycan types on the sequons of N74, N331, N343, and N801 did not change between two spike proteins, although some Omicron mutations, including del69-70, G339D, and D796Y, occurred only a few residues away upstream or downstream of these N-glycosylation sites. This suggests that the glycosylation processing on these sites was not altered by any structural changes caused by nearby mutations on S-Omicron.

An unusual situation was observed for the glycosylation at the N149 site during MS based analysis of glycopeptides derived from SARS-CoV-2 spike proteins. N149 is a surface residue in the NTD of the spikes (Fig. 1) and is within the area of antibody binding NTD supersite (*51*). Multiple residues representing specific cleavage sites for certain common proteases such as trypsin (K/R), chymotrypsin (F/Y/W/L), and alphalytic proteases (A/V/S/T) are in the sequences flanking N149 (Fig. S2A). This implies it should be easy to produce detectable N149-containing glycopeptides by various enzymes or protease combinations. However, no high quality data were produced for quantifying the glycan abundance at this site from the analysis of D614G, Alpha, Beta, Gamma and Delta spike variants in our previous study (*37, 38*) and this phenomenon also has been observed by other laboratories (*30*). In this report the sequential digestion by Asp-N and chymotrypsin under selected condition allowed quantitative characterization of N149 glycosylation on S-Omicron and S-D614G with the production of glycopeptides of DHKNN(glycans)KSWMESEF and YHKNN(glycans)KSWMESEF, respectively (Fig. S2A). On the other hand, from the digestions by Lys-C /Lys-C, Lys-C / trypsin, and Lys-C/ alpha-lytic protease, a comparable number of the N-glycans at this site were determined for S-Omicron. However, only one glycan was detected from D-614G (Table S1), suggesting that mutations of the G142D and del143-145 on S-Omicron (Fig. 1) might lead to the formation of more detectable glycopeptides from this protein than from D614G and other variants of concern.

To further understand the role of the N-glycosylation at N149 on the interaction of the spike protein with receptor and antibody, we prepared two spike mutants, S-Omicron-N149Q and S-D614G-N149Q, and evaluated their binding capability to a human ACE2 and a monoclonal antibody (4A8) against the NTD of spike protein. 1D gel analysis of pull-down experiments showed that comparable amounts of four proteins, either wild type S proteins or N149Q mutants, were able to bind to Fc tagged ACE2 proteins that were immobilized on protein G magnetic beads (Fig. S2B, lanes 5, 8, 11, and 14), revealing that the carbohydrate groups at this site were not essential for the binding between S and the receptor proteins. We observed that 4A8-bound S-D614G-N149Q increased by approximately 25% in comparison with S-D614G (Fig. S2C, lanes 2 and 4), suggesting the potential impairment by this glycan-modified residue on antibody recognition. Because the residues of H146 and Y145 of S protein have direct contact with mAb residues and glycosylated N149 is close to the complex interface revealed by the Cryo-EM structure of the S protein-4A8 complex (*52*), increased binding of S-D614G-N149Q to 4A8 suggests that the N-glycans on N149 might participate the interactions between two molecules. On the other hand, no binding of wildtype and mutated S-Omicron proteins to the 4A8 mAb was observed, presumably due to the loss of contact residues because of variant-specific mutations in S-Omicron, including deletion of residues V143 to Y145 and/or substitution of G142D. Further investigation is needed to understand the role of N149 glycosylation in this Omicron variant.

### Analysis of disulfide bonds

Fifteen disulfide bonds formed by 30 cysteine residues in the ectodomain of the S proteins have been visualized by three-dimensional structures and mass spectrometry (*13, 40*).

Although these disulfide bonds are conserved in all variants of concerns, Yao et al. have revealed that some disulfides in the RBD of Omicron S protein, such as C480-C488 and C379-C432, are susceptible to reduction that could affect binding capacity and stability of the protein(*49*). To understand whether mutations of Omicron S affect the formation of disulfide bonds, we examined and compared the disulfide bonds of the S proteins of Omicron with the D614G variant using mass spectrometry analysis with four various enzyme digestion methods.

As depicted in Fig. S3, the peptides containing disulfide bonds and free cysteines could be unambiguously detected through HCD-induced fragmentation of protein digests. This approach allowed the detection of all disulfide bonds, except C15-C136, C131-C166, and C617-649, with one or more interpeptide or intrapeptide disulfide-bond containing peptides for each bond (Table S1). Fig. 4 shows that the abundance of disulfide-bond related peptides on Omicron and D614G S proteins were well correlated. It suggests that these spike proteins possess similar overall disulfide bond structures and that the large number of variant-specific mutations in Omicron S might not lead to a significant alteration on its disulfide bond linkages. Additionally, peptides with free cysteine modified by NME were detected on most of the cysteine residues (data not shown), indicating that a fraction of each of these residues does not form proper disulfide bonds before sample preparation. This could be because of incomplete formation or partial reduction of these disulfide bonds during protein expression or storage. However, accurate quantification of individual disulfide bonds was limited by relatively high variations for some disulfide bonds represented by multiple peptides (Table S1). This presumbly would be caused by more nonspecific cleavage for disulfide-bonded peptides than for linear peptides during enzyme digestion, because four cleavages for each inter-peptide disulfide bonded molecule are required and the spatial structure of such di-peptides may also hinder the effective access of proteases. More optimization of sample preparation and the digestion procedure is needed to generate uniform disulfide-bonded peptides for more accurate disulfide bond quantification.

**Fig. 4.**
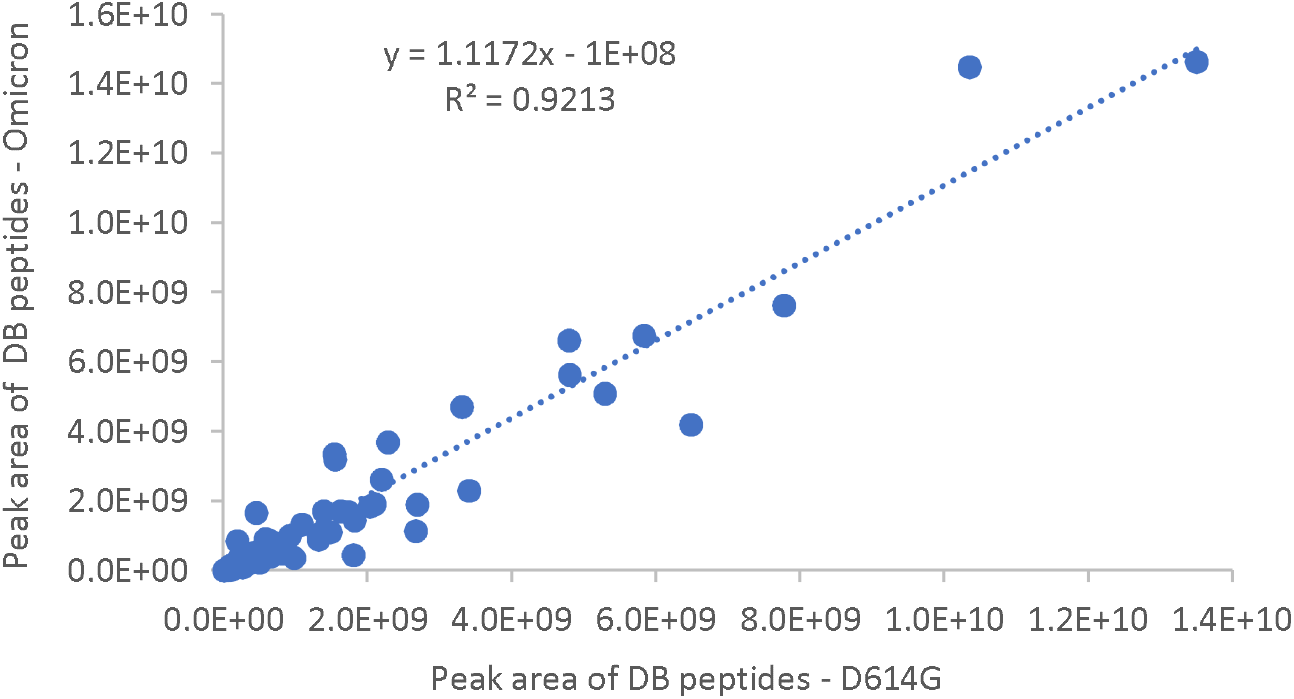
Correlation of the disulfide-bond containing peptides detected from SARS-CoV-2 spike proteins of the Omicron and D614G variants.

### Molecular dynamic simulations

Glycans, vital components of the S protein, fulfill various key functions, including stabilizing the protein structure and promoting immune evasion(*23, 53-56*). According to the latest update (June 13, 2023) from the CoV-AbDab (Coronavirus Antibody Database)(*57*), of the total 12,536 antibody entries, nearly all (12,431) target the S protein, signifying its central role in antibody-based interventions. Glycosylation frequently aids the virus in circumventing the immune system by creating a shield over critical epitopes on the S protein, thereby reducing the antibodies’ effectiveness.

To understand how the differences in the glycosylation profiles of the Omicron and D614G variants affect glycan shielding and other roles of glycans, we built all-atom models for both variants’ S proteins, incorporating glycans (Fig. S5). The selection and construction of specific glycans integrated into these models were guided by glycosylation profile data derived from mass spectrometry analysis detailed above. Subsequently, we carried out three 1.4-µs molecular dynamics simulations for each system; we also simulated the same protein models with no glycans present. This comprehensive approach allows us to resolve the role of glycosylation in viral immune evasion strategies and can help guide the development of more effective antibody-based interventions against different SARS-CoV-2 variants.

Inspired by the work of Casalino et al.(*23*), we provide an overall view of the S protein’s glycan shield (Fig. 5). Fig. 5A and 5B shows the superimposition of glycans over the course of the simulations, demonstrating the range of potential conformations. Despite the use of smaller intervals in previous studies(*23*), this representation still offers a realistic depiction of the glycan shielding. Given that the process of antibody binding takes place over microseconds, the chosen interval of 0.25 μs strikes a suitable balance between computational feasibility and the accuracy of the model.

**Fig. 5.**
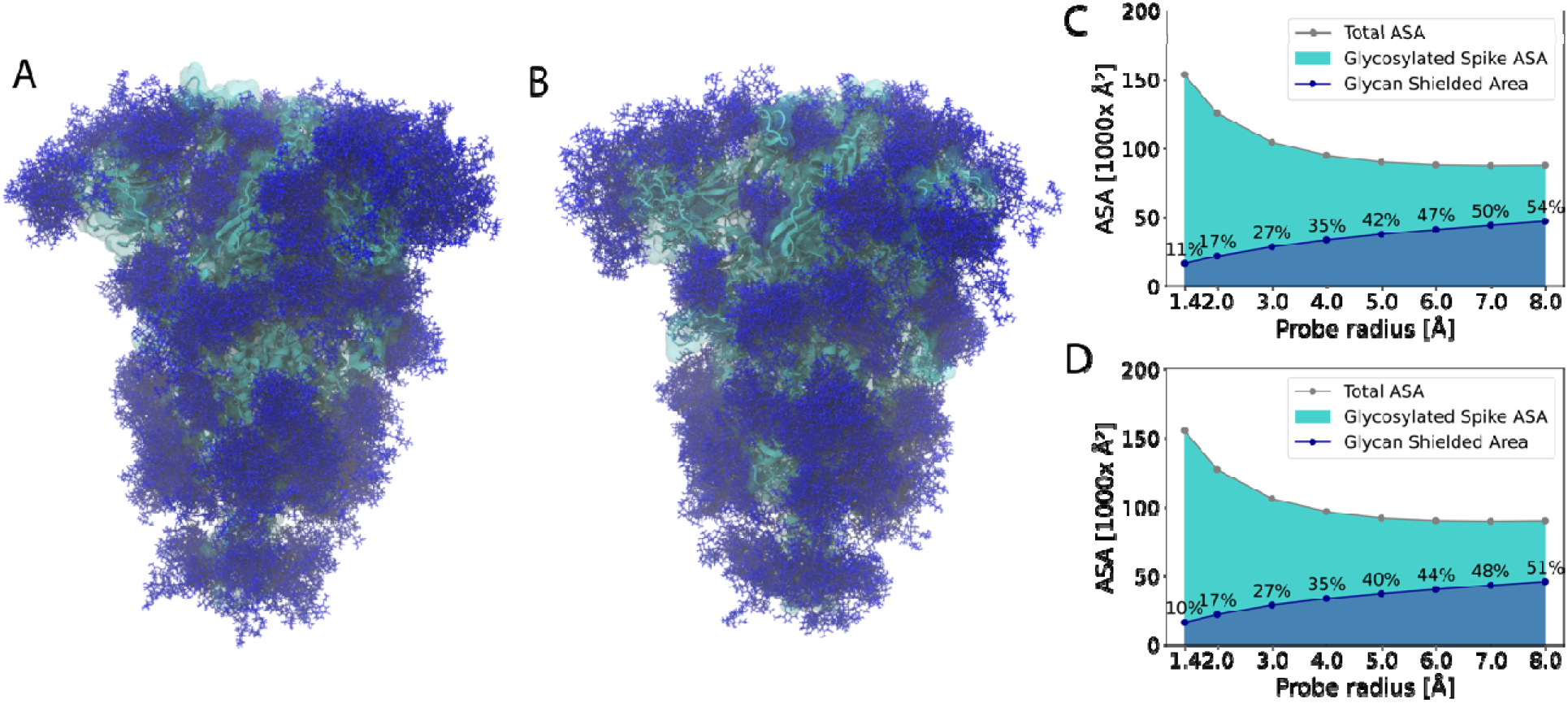
Glycan shield of the SARS-CoV-2 S protein. The D614G (**A**) and Omicron (**B**) S proteins are depicted in cyan using a cartoon representation overlaid with a transparent surface. The superimposed glycans are represented as dark blue sticks. These glycan conformations were captured at intervals of 0.25 μs throughout the net 4.2 μs of simulation trajectories (3 × 1.4 μs) for the D614G and Omicron S proteins. The accessible surface area (ASA) of D614G (**C**) and Omicron (**D**) S proteins was evaluated using a variety of probe sizes, spanning from 1.4 Å to 8 Å.

To comprehensively estimate the accessibility of S proteins under the glycan shield and identify the differences between the two variants, we performed accessible surface area (ASA) calculations using probe radii ranging from 1.4 Å to 8 Å. These different probe radii allow us to estimate accessibility for molecules of different dimensions —from smaller ones, such as water, to larger entities, such as antibodies. The 1.4-Å probe radius is typically used to mimic water’s accessibility. In assessing the accessible surface area for antibodies, we employed larger probe radii, reaching up to 8 Å. The probe radii are proven to be effective when compared with the 5 Å to 10 Å range used in previous research (*20, 56, 58, 59*). We observed that even without glycans, larger molecules have less accessibility to the S proteins than do smaller ones (Fig. 5C and 5D). Furthermore, the presence of glycans exacerbates this, making it increasingly difficult for larger molecules to access the S proteins. However, what is most striking is that when comparing the Omicron and D614G S proteins, the Omicron variant exhibits a notably greater ASA for large molecules.

Among the various epitopes present on the S protein of SARS-CoV-2, the RBD is notably the most targeted by antibodies(*60, 61*). In fact, according to the CoV-AbDab, out of the 12,431 antibodies that target the S protein, 8,393 are specifically directed towards the RBD(*57*). To gain further insights into the epitopes of the RBD, we specifically examined the glycan shield present on the RBD (Fig. 6A-D). Intriguingly, our observations reveal that the RBD in chain A is enveloped by numerous glycans originating from chain B (Fig. 6A,B). This pattern of glycan shielding is also evident when examining the RBDs in chains B and C, further illuminating the complex interplay of these structures in the virus’s immune evasion strategies.

**Fig. 6.**
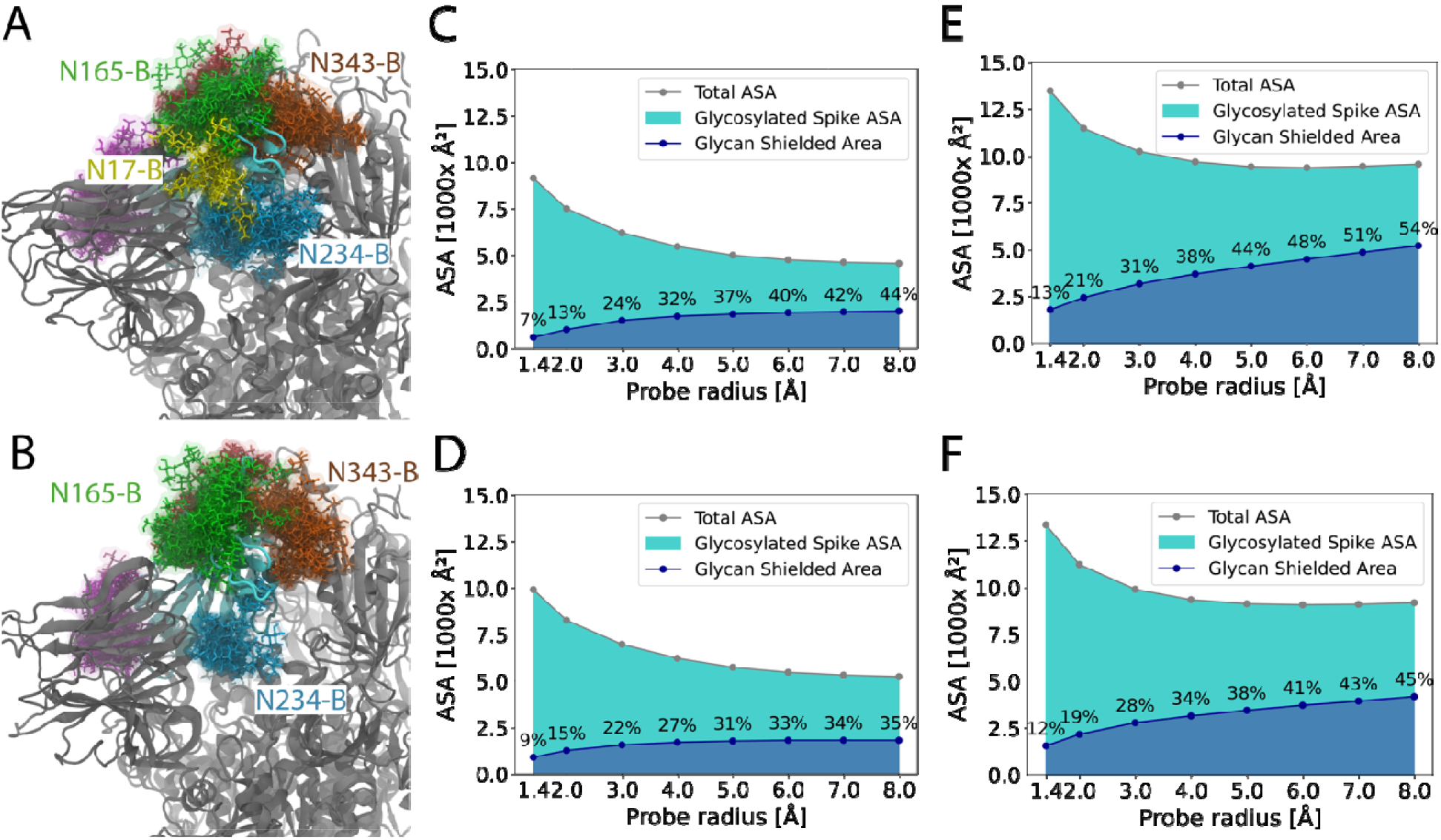
Glycan shielding of the receptor binding domain (RBD) and N-terminal domains (NTD) in the SARS-CoV-2 S protein. The RBDs of chain A in the D614G (**A**) and Omicron (**B**) S proteins are shown in cyan. All glycan residues within 5 Å of the RBDs are depicted by differently colored stick representations. These glycan structures are superimposed at intervals of 0.25 μs along the respective simulation trajectories. The ASA of RBDs in the D614G (**C**) and Omicron (**D**) S proteins was evaluated using a variety of probe sizes, spanning from 1.4 Å to 8 Å. The ASA of NTDs was also evaluated for both D614G © and Omicron (F) S proteins.

In the case of the D614G variant, the glycan at position N17 from chain B wraps around the RBD, which is not observed in the Omicron variant. The most abundant glycan at position N17 is H2H4F1 in Omicron and N5H4A1 in D614G. This results in a shorter N17 glycan in Omicron than in D614G, which is likely why the N17 glycan wraps around the RBD in D614G but not in Omicron. The absence of the N17 glycan around the RBD in Omicron results in residues from Ser469 to Val483 having a larger accessible area in this variant (Fig. 6A, B, S6). It is noteworthy that the glycosylation site at N17 in the Delta variant is absent, attributable to the T19R mutation. An absence of glycans at the N17 position in the Gamma variant’s S protein also has been noted previously(*33, 62*).

To further identify potential epitopes within the RBD, we analyzed the ASA of each residue, taking into account the presence of glycans (Fig. S8). In this analysis, residues ranging from Ser469 to Val483 in the Omicron variant again exhibited a larger ASA. Also noteworthy, the S477N and T478K mutations in the Omicron variant further amplify the ASA of the RBD in its S protein. The combination of a shorter or absent glycan at N17, along with protein mutations, makes this region a promising potential epitope. In a broader perspective, when comparing the RBDs in the Omicron and the D614G variants, the glycan shield over the RBD is less extensive in the Omicron variant than in D614G, especially for larger molecules (Fig. 6C,D). Specifically, for molecules with a radius of 8 Å, the Omicron variant RBD has 35% of the area shielded by glycans, whereas the D614G variant has 44% of its RBD area shielded.

Besides the RBD, another potential epitope is the N-terminal domain (NTD). The NTD in the Omicron variant is less shielded by glycans than is the NTD in the D614G variant (Fig. 6). In particular, for molecules with a radius of 8 Å, the Omicron variant has 45% of the area covered by glycans, whereas the D614G variant has 54% of its area shielded by glycans. The most frequently observed glycan at position N149 is N2H5 in Omicron and N4H5F1A1 in D614G, respectively. This results in a shorter N149 glycan in Omicron than in D614G. Thus, changes in the glycosylation profile at N149 and N17 contribute to a smaller glycan shield on the NTD of Omicron. Fig. S9 shows the extent of glycan shielding of the NTDs. Fig. S10 shows the full-length glycans at positions N17 and N149. We observed that the secondary structure in the Omicron variant, specifically two β-strands and a loop between them spanning from residue 140 to 158, exhibited decreased stability. This instability might be attributed to the deletion of residues V143, Y144 and Y145, as well as the G142D mutation in this region.

In addition to promoting immune evasion, glycans play a pivotal role in stabilizing the S-protein structure. In two out of the three 1.4-μs simulations of the Omicron S protein without glycans, which started in a down (closed) conformation, we identified a “sub-down” RBD conformation (Fig. 7A). This sub-down RBD conformation does not manifest in any simulations conducted in the presence of glycans (Fig. 7C). To quantify the RBD conformations, we introduced two collective variables, distance and dihedral angle, consistent with definitions used previously(*56*). We used two-dimensional KDE plots to visually represent the distribution of these two variables concurrently (Fig. 7B,D). To provide context, an “up” or open conformation of the RBD is defined by a distance of 70 Å and a dihedral angle of 0°. In the simulations devoid of glycans, we discerned two clusters, one representing the “down” or closed RBD conformation and the other depicting the sub-down conformation (Fig. 7B). Both conformations are shown in Fig. 7A. In contrast, the distribution of RBD conformations in the presence of glycans, as seen in all three replicas, is exclusively located within the down or closed-state cluster. Furthermore, it appears more intense and concentrated around the distribution center, indicating that the presence of glycans stabilizes the RBD conformation. Additionally, we observed glycans N165, N234, and N343 enveloping the RBD (Fig. 7C), in agreement with previous observations(*56*). This further supports the idea that the glycans surrounding the RBD stabilize it, preventing the emergence of a sub-down conformation.

**Fig. 7.**
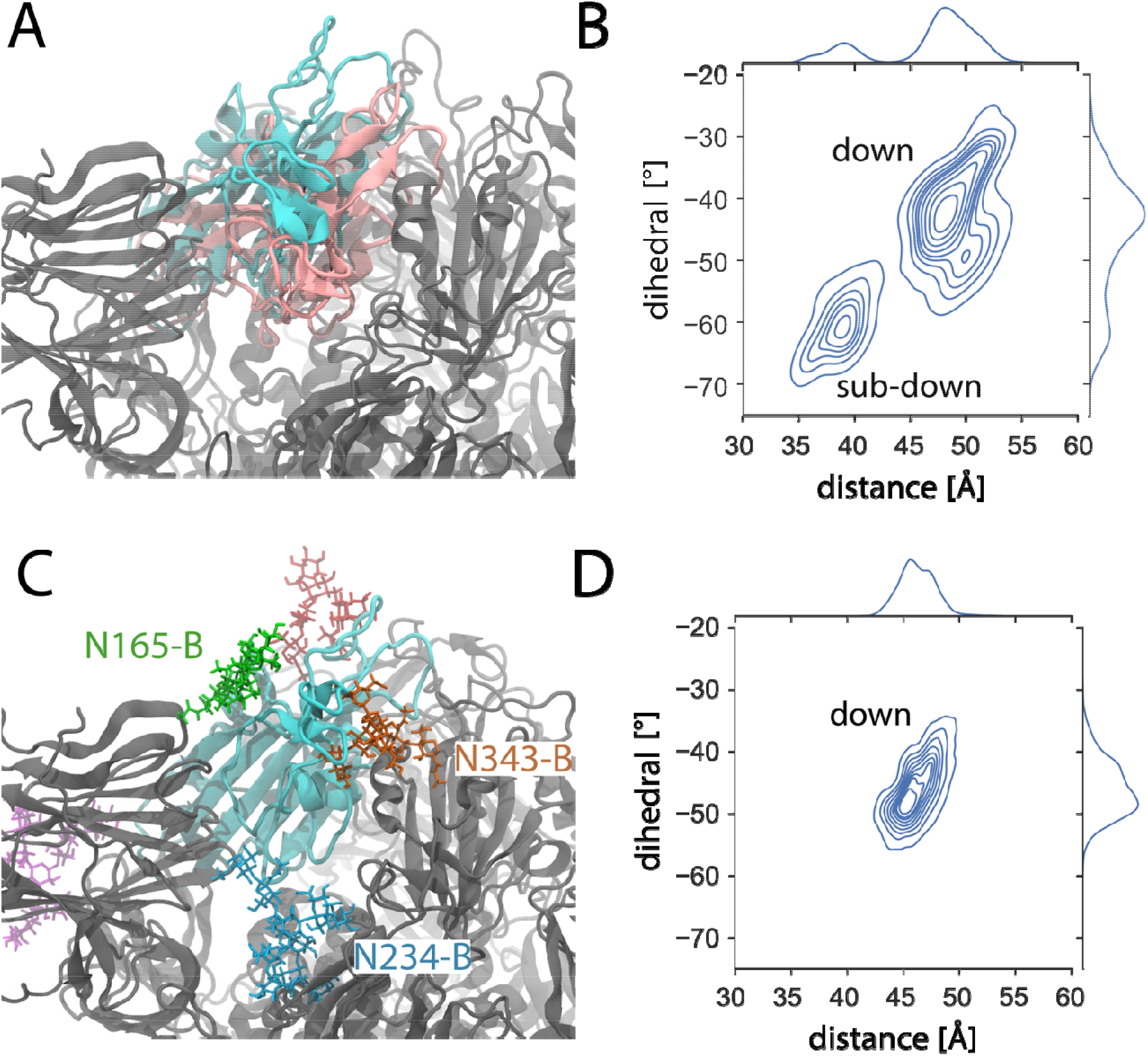
Sub-down RBD conformation in the Omicron S protein in the absence of glycans. (**A**) Initial RBD conformation (PDB: 7TNW(*63*)) and sub-down RBD conformation are depicted in cyan and pink, respectively, in the simulations conducted without glycans. (**B**) A two-dimensional kernel density estimate (KDE) plot visualizes the spread of RBD conformations within the trajectories of the un-glycosylated systems, with two collective variables used previously(*56*) (defined in Methods), a distance and a dihedral angle, characterizing the RBD conformations. (**C**) The RBD conformation in simulations that incorporate glycans. Glycans surrounding the RBD are depicted in differently colored stick representations. (**D**) The corresponding KDE plot illustrates the distribution of RBD conformations in the glycosylated simulations.

### Correlation of MD simulations and hydrogen/deuterium exchange mass spectrometry

One quantity that can be extracted from MD simulations is the root mean squared fluctuation (RMSF) of each residue, which is an estimate of the residue’s flexibility during the simulation. Hydrogen/deuterium exchange mass spectrometry (HDX-MS) measures changes in a protein’s D_2_O uptake, which reflects changes in hydrogen bond stability. The implication of both techniques is that changes in the protein’s conformational dynamics are being detected, e.g., high RMSF values may correlate with high percentage deuteration. We tested this hypothesis by comparing RMSF and HDX-MS data per residue for the D614G variant with an x-y scatterplot and filtering for RMSF > 4 Å and deuteration > 5%. We note that some regions of high deuteration did not correspond to high RMSF, possibly due to the disparate timescales explored by the methods (microseconds for MD and minutes for HDX-MS). Nonetheless, 6 regions satisfied both criteria: residues 72-75, 147-150, 248-256, 445-447, 475-486, and 677-688 (Fig. 8A). The first three regions are adjacent to each other in the NTD, the next two regions are the receptor binding motif (RBM) of the RBD, and the last region includes the furin cleavage site between residues 685 and 686. Visualizing these six regions on the D614G S protein structure (6VSB, Fig. 8B) indicates they are all on the surface, and the key roles of the RBMs and furin cleavage site in SARS-CoV-2 infection are well-documented(*12, 13*). Therefore, the correlation of RMSF in MD simulations and HDX-MS data indicates that the S protein’s more dynamic regions also have important biological functions.

**Fig. 8.**
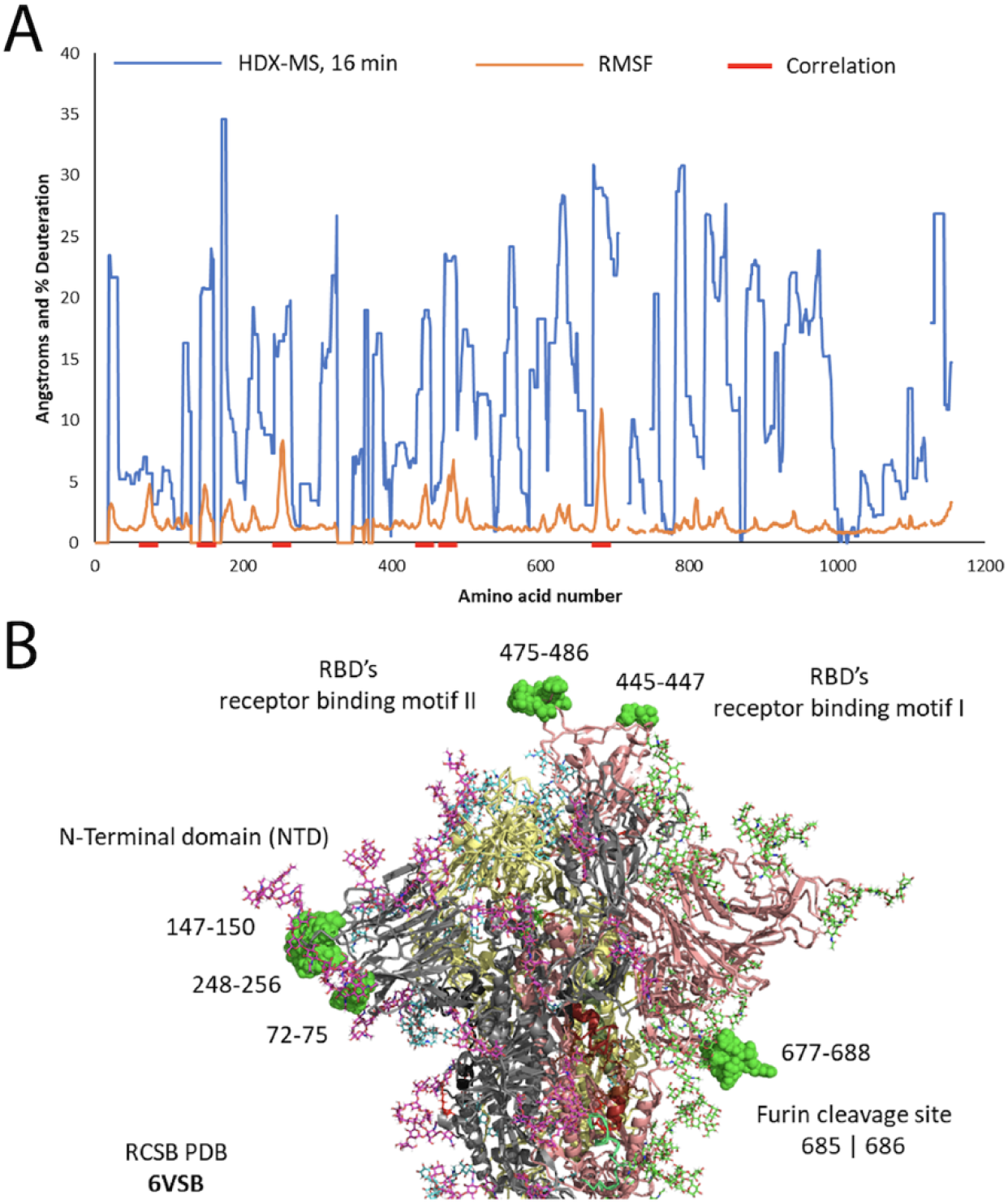
(**A**) Root Mean Square Fluctuations (RMSF) and percentage of Hydrogen/Deuterium Exchange Mass Spectrometry (HDX-MS) at 960 sec for each residue in the S protein. The RMSF, obtained from MD simulations, shows the average level of fluctuations in the protein’s structure across different chains and replicas. Gaps in HDX-MS sequence coverage are shown as breaks in both lines. Correlation of the two data types and filtering for RMSF > 4 Å and deuteration > 5% selected the regions shown as red bars: residues 72-75, 147-150, 248-256, 445-447, 475-486, and 677-688. (**B**) Model of S glycoprotein trimer (PDB 6VSB) with glycans and correlations between HDX-MS and MD RMSF. The S protein monomers are grey, yellow, and salmon with magenta, blue and green glycans, respectively. The six regions correlated by the highest HDX-MS and MD RMSF measurements are shown as green space-filling residues on the NTD (72-75, 147-150 and 248-256), the RBD (445-447 and 475-486) and the furin cleavage site between 685 and 686 (677-688).

## Discussion

In this comprehensive study, we used advanced LC-MS techniques to conduct an analysis of the N-glycosylation and disulfide bond profiles of the recombinant spike ectodomain protein derived from the Omicron variant, while using the D614G S protein as a control. To ensure the accuracy and validity of our results, we used the proteins expressed and purified under identical experimental conditions and selected N-glycosylation data only when multiple datasets from distinct digestion experiments showed consistent outcomes. Through these strategies, we determined the distribution and relative abundance across all 22 potential N-glycosylation sites of the S-Omicron and S-D614G proteins. We also detected disulfide bonds within the two proteins. Our data reveal that the Omicron S protein aligns closely with the glycan processing patterns of S-D614G at the majority of sequons. Meanwhile, significant differences in glycan types were observed at specific sites, namely N61, N149, N657 and N1098. In terms of the predominant glycan abundances at various sequons, distinctions were evident at multiple sites, including N17, N149, N331, and N657. Our investigation also detected 12 out of 15 disulfide bonds, with close similarity between two proteins, by comparing the intensities of disulfide-formed and free sulfur-containing peptides. Moreover, we conducted MD simulations of both protein variants, with and without the inclusion of major glycans, shedding light on the effects of N-glycosylation on protein structure. Our simulations revealed that S-Omicron showed diminished glycan shielding in comparison to S-D614G. The difference was particularly pronounced within the RBD and NTD regions, which encompass potential epitope areas. These findings enhance our understanding of the intricate interplay between glycosylation, protein structure, and immune evasion strategies. As the global pursuit of effective vaccination and therapeutic strategies continues, our research contributes vital insights to this ongoing endeavor.

## Materials and Methods

### Materials

All chemicals were obtained from Sigma–Aldrich (St. Louis, MO) except where otherwise indicated. Endoproteases including trypsin, chymotrypsin, Lys-C, and Asp-N were purchased from Promega (Madison, WI), and a-lytic protease was obtained from New England BioLabs (Ipswich, MA). The sequences of the proteins include the mutated furin cleavage site and six proline substitutions at F817P, A892P, A899P, A942P, K986P, and V987P to stabilize the prefusion conformation of the trimeric spike proteins. Phosphate-buffered saline (PBS) tablets, LC-MS grade water, 0.1% formic acid, methanol, and acetonitrile 0.1% formic acid were from Fisher Scientific (PA). Deuterium oxide (99.9%) was from Cambridge Isotope Laboratories (MA). Urea and tris(carboxyethyl)phosphine (TCEP) were from Sigma-Aldrich (MO). Reagent and sample vials for HDX-MS were from Trajan Scientific and Medical (NC) and Thermo Scientific (CA), respectively.

### Preparation of recombinant spike proteins and analysis workflow

The recombinant ectodomains of the spike proteins of SARS-CoV-2 Omicron and D614G variants were prepared in HEK cells under identical expression and purification conditions. In particular, The poly-histidine-tagged and six-proline-stabilized ectodomains of the SARS-CoV-2 S proteins including S-Omicron, S-D614G, S-Omicron-N149Q, and S-D614G-N149Q were expressed in human Expi293F cells using the Expi293 Expression System (ThermoFisher Scientific) and purified using a HisTrap FF column (GE Life Sciences)(*64*) followed by gel filtration on a Superose 6 Increase 10/300 GL column (GE Life Sciences). Alkylation, reduction, and digestion of the samples were conducted at pH 7.9 for N-glycosylation analysis. Sample preparation for detection of disulfide bonds was performed at pH 6.5 to reduce the formation of scrambled disulfide bonds (Fig. S1). The proteins also were treated with N-ethylmaleimide (NEM) to permanently block sulfhydryls of free cysteine residues to prevent them from oxidation and formation of artificial disulfide bonds. The most abundant N-glycan at each sequon of the two S proteins obtained from N-glycosylation analysis was incorporated into the protein models for MD simulations.

### Digestion of spike proteins

The enzymatic digestion of recombinant proteins was conducted as previously reported (*37*). In brief, aliquots of 1-2 μg full-length ectodomain spike protein were denatured and reduced at 60 °C for 30 min in a solution containing 50 mM ammonium bicarbonate (pH 7.9), 0.05% RapiGest SF surfactant (Waters Corporation) and 5 mM DTT. Samples were alkylated using 15 mM iodoacetamide for 30 min in the dark at room temperature with gentle mixing. Digestions with a single enzyme were conducted at 37°C overnight at an enzyme-to-protein ratio of 1:10 (w/w). For the samples digested sequentially by two enzymes, the first digestion was performed at 52 °C for 60 min with the first protease (1:3 w/w). The second digestion was conducted at 37 °C overnight at an enzyme-to-substrate ratio of 1:15 (w/w). The proteolytic reactions were quenched and the RapiGest was precipitated by adding 5% trifluoroacetic acid to decrease the pH to below 3. The mixture was then incubated at 37 °C for 30 min. The solutions were centrifuged at 4000 RPM for 10 min, and the supernatants (20 μL) were transferred into new sample vials. Each digestion was performed with three replicates.

For the characterization of disulfide bonds, proteins were alkylated by 1 mM Pierce N-ethylmaleimide (ThermoFisher Scientific) in alkylation buffer containing 8M urea, 100 mM Tris (pH 6.5) at 37 °C for 2 hours. The pretreated proteins were then precipitated by adding chilled acetone to the reaction solutions, and the pellets were resuspended with 100 mM ammonium citrate (pH 6.5) followed by sequential digestion.

### Mass spectrometry analysis

LC-MS analysis of protein digests was performed using an Orbitrap Eclipse Tribrid mass spectrometer coupled to an UltiMate3000 RSLCnano chromatography system (Thermo Scientific) as described previously(*37*). The peptides were injected into an integrated separation column/nanospray device (Thermo Scientific EASY-Spray PepMap RSLC C_18_, 75 µm i.d. × 15 cm length, 3µm 100Å particles), coupled to an EASY-Spray ion source. Mobile phase A (0.1% formic acid) and mobile phase B (0.1% formic acid in 80% acetonitrile) were mixed based on the following gradient with the flow rate of 300 nL/min: 4% B for 8 min; 4 – 10% B for 2 min; 10 – 35% B for 33 min; 35 – 60% B for 2 min; and 60 – 95% B in 1 min. The spray voltage was set to 1.8 kV, and the temperatures of the integrated column/nanospray device and the ion transfer tube were set at 55 °C and 275 °C, respectively.

MS data acquisition was accomplished in positive ion mode using a signature ion triggered electron-transfer/higher-energy collision dissociation (EThcD) method. The full MS precursor scans were acquired by the Orbitrap at a resolution of 120,000 at *m/z* 200, from *m/z* 375 – 2000 with the automatic gain control (AGC) target setting as “standard” and the maximum injection time as “auto”. After the MS1 survey scan, a data-dependent MS2 scan was acquired over a 3-s cycle time using high-energy collision dissociation (HCD) at a resolution of 30,000, a mass range of *m/z* 120 – 2000, and a normalized collision energy (NCE) of 28%. Signature ions representing glycan oxonium fragments were used to trigger the electron-transfer dissociation (ETD) fragmentation. If one of three common glycan signature ions—*m/z* 204.0867 (HexNAc), 138.0545 (HexNAc fragment), or 366.1396 (HexNAcHex) —was detected in the HCD spectrum within a 15 ppm mass accuracy, an additional precursor isolation and EThcD acquisition were performed. Settings for that included a resolution of 50,000 at *m/z* 200, with the range of *m/z* 150 – 2000, 500% normalized AGC target, 150 ms maximum injection time, and 35% supplemental activation NCE.

### Data analysis

MS/MS data for the analysis of glycopeptides were processed using the PMi-Byonic (version 3.7) node within Proteome Discover (ThermoFisher Scientific). Data were searched using the Protein Metrics 182 human N-glycan library (included in the Byonic program) as potential glycan modifications. The search parameters for enzyme digestion were set to semi-specific, three allowed missed cleavage sites, and 6 ppm and 20 ppm mass tolerance for precursors and fragment ions, respectively. Carbamidomethylation of cysteine was set as a fixed modification with variable modifications set to include deamidation at Asn and Gln and oxidation of Met. MS2 spectra of identified glycopeptides with a Byonic score higher than 150 were considered valid identifications. Identified glycopeptide and unoccupied peptide abundances were determined using precursor ion peak intensity with normalization on total peptide amount per file. Relative abundance of each type of glycans at each site was calculated as the normalized peak intensity ratio of the peptides bearing a particular glycan type over sum of total glycopeptide intensity. The glycan abundance was represented as the mean of three replicates along with standard deviation of the mean. Data for disulfide bonded peptides were processed using PMi-Byos (Protein Metrics). The search parameters for enzyme digestion of disulfide bonded peptides in Byos were set to fully-specific, four allowed missed cleavage sites, and 6 ppm and 15 ppm mass tolerance for precursors and fragment ions, respectively. Oxidation of Met and Trp, deamidation of Asn, and thioether formation by derivatization with NEM on Cys, were considered as variable modifications. N-glycans were not included in the disulfide search due to increased complexity of data generated. Peptide quantitation was performed using the Byologic module with fully specific digestion and at specific sites corresponding to each digestion enzyme combination.

### Binding assay and 1D Gel electrophoresis

The binding of S proteins to human ACE2 and monoclonal antibody (mAb) was carried out on magnetic beads. The Fc-tagged ACE2 or mAb were first immobilized on Protein G Dynabeads (ThermoFisher Scientific). The S protein solutions were then incubated with on-bead receptor or antibody at room temperature for 60 min. Bound S protein/ACE2 or S protein/mAb complexes were analyzed on sodium dodecyl sulfate–polyacrylamide gel electrophoresis (SDS-PAGE) on a NuPAGE Novex Bis-Tris gel following manufacturer’s instructions (Invitrogen). Protein solutions were mixed with 4× sample buffer and deionized water (1:3 v/v) and heated at 80 °C for 10 min. The supernatants were loaded on a 4–12% gradient gel, and the gel was run in MOPS buffer at 200 V for 45 min. The gels were stained with the Invitrogen SYPRO Ruby protein gel stain (ThermoFisher Scientific).

### Hydrogen/Deuterium Exchange Mass Spectrometry

A DHR-PAL system (Trajan) was used for sample preparation, with sample tray at 20 °C, quench tray at 4 °C, valve chamber & pre-chiller at 4 °C, and digestion chamber at 8 °C. Purified spike ectodomain (0.4 μg/μL) was mixed 1:5 (v/v) with PBS prepared using H_2_O (equilibration buffer, measured pH 7.22) or D_2_O (labeling buffer, measured pH 7.56). Labeling times were 0, 60, 240, and 960 sec with 3 technical replicates at each time. Samples were quenched with an equal volume of 2 M urea 0.5 M TCEP pH 2.5 and held at 4 °C for 2 min. An UltiMate3000 UPLC system (binary nano pump and loading pump, Thermo Scientific) was used for subsequent online sample handling. Automated valve switching passed the quenched sample over a 2.1 × 20 mm Nepenthesin-2 / Pepsin mixed digestion column (AffiPro, CZ) at 100 μL/min H_2_O 0.1% HCOOH for 2 minutes, trapping the resulting peptides on a 2.1 × 5 mm Fully Porous C18 guard column (Phenomenex, CA), then de-salted peptides at 300 μL/min for 4 minutes. Peptides were eluted and resolved by a gradient from 13 to 65% mobile phase B (95:5:0.1 CH_3_CN / H_2_O / HCOOH) over 23 min on a 1 × 100 mm Luna Omega 1.6 μm 100 Å C18 column (Phenomenex, CA). A Tribrid Eclipse Orbitrap (OT) mass spectrometer (Thermo Scientific) with HESI-2 electrospray ion source and high-flow needle was operated in positive ion mode to detect peptides for 31 minutes. In all samples precursor scans of resolution 120K (at *m/z* 200) in the range 375-2000 *m/z* were acquired. For each MS1 scan, the top-10 abundant precursor features were selected for data-dependent MS2 scans, selecting for an intensity threshold of 30K counts, monoisotopic peptide precursors, charge states 2^+^ to 8^+^ with an isolation window of 1.2 *m/z*, and not repeating precursor ions more than twice within 15 sec. Precursors were fragmented with HCD at 28% normalized collision energy (NCE) and centroid scanned in the OT with standard automatic gain control (AGC) target, automatic injection time, scan range 120-2000 *m/z*, and resolution 30K. If at least one of three selected oxonium ions was detected (HexNAc 204.0867, HexNAc fragment 138.0545, or HexNAcHex 366.1396) with 15 ppm mass tolerance, then EThcD OT-MS2 scans were acquired of that precursor ion. Supplemental HCD was at 20% NCE with profile scans from 150-2000 *m/z* at resolution 50K using custom AGC target (500%) and fill time (90 msec). At the end of the analytical gradient, solvent transitioned to 90% mobile phase B for 6 min, and halfway through that time the analytical and trapping columns were put into back-flow washing mode by automated valve switching. After re-equilibration of the analytical column at 13% mobile phase B the injection cycle ended at 45 min. As recommended(*65*), the entire batch of control and heat denatured samples (all time points and technical replicates) was randomized for HDX-MS acquisition to minimize batch effects on interpreted differences in protein state, labeling time, and replicates.

### HDX-MS Data Processing

Protein Metrics Inc. (CA) Byos HDX 4.6-37 searched the 3 data files from equilibration buffer samples (0 sec labeling time, 3 technical replicates) to identify (glyco)peptides using MS2 spectra. The search database included spike and both proteases. Both HCD and EThcD tandem mass spectra contributed to peptide spectral matching. Putative (glyco)peptides were then searched for in data files from all samples at the MS level (and appropriate retention times) to identify both unlabeled and deuterated peptides and visualize their isotopic envelopes. Initial spike results (1271 peptides) were narrowed by 1) default software filters (MS2 score > 15, minimum alt_rank_score/primary_rank_score > 0.99, maximum precursor *m/z* error ± 40 ppm, maximum retention time deviation ± 5 min) leaving 1056 peptides, and 2) removing peptides with MS2 score < 150 leaving 106 of 205 glycopeptides, removing peptides with more than ± 10% average, maximum, or minimum “deuteration” in 0 sec samples, and removing peptides causing standard deviations >10% at any labeled time-point, leaving 561 peptides. Additional manual curation involved adjustment of the extracted ion chromatogram (XIC) window used to integrate MS data and generate an isotopic envelope, optimizing the intensity and specificity of that envelope. Peptides with inadequate intensity XICs to estimate deuteration were discarded.

### Molecular dynamic simulations

We selected S-protein structures for simulation by aiming for those at the highest resolution and with the fewest artificial mutations. The wild type (WT) Omicron S protein was initialized from the structure in the RCSB Protein Data Bank (PDB) ID: 7TNW(*63*). Missing residues in 7TNW from 245 to 247 and 255 to 259 were copied from PDB ID: 7WK2(*66*). Residue 141 was mutated to leucine, according to the sequence of Omicron S protein used in this study. On the basis of its local electrostatic environment, His625 was set as HSD (histidine delta; protonated on its δ nitrogen); all other histidines were set as HSE (histidine epsilon; protonated on the ε nitrogen). Using the WT S protein as a starting point, we also built the common laboratory version of the Omicron S protein. Here, residues 682-685 were mutated to the sequence GSAS, and residues 817, 892, 899, 942, 986, and 987 were mutated to proline. The lab version of D614G S protein was initialized from the PDB ID:7KRQ(*67*). Based on its local electrostatic environment, His49 was set as HSD and all other His were set as HSE. Residues 817, 892, 899, 942, 986, and 987 were mutated to proline.

Disulfide bonds between residues Cys15 and Cys136, Cys131 and Cys166, Cys291 and Cys301, Cys336 and Cys361, Cys379 and Cys432, Cys391 and Cys525, Cys480 and Cys488, Cys538 and Cys590, Cys617 and Cys649, Cys662 and Cys671, Cys738 and Cys760, Cys743 and Cys749, Cys840 and Cys851, Cys1032 and Cys1043, and Cys1082 and Cys1126 were added in all systems.

Glycosylation sites are located on N17, N61, N74, N122, N149, N165, N234, N282, N331, N343, N603, N616, N657, N709, N717, N801, N1074, N1098, N1134, and N1158 in all glycosylated systems. Overall, 20 N-linked glycans are present in each protomer, resulting in a total of 60 glycans for one S-protein trimer model. The glycan at each site with the highest population in the mass spectroscopy data (Fig. 4) was added to the site. Among the possible glycan structures with the same constitution according to mass spectroscopy, we selected the most populous one with the highest "hit score" from the GlyGen website (https://www.glygen.org/glycan-search)(*68*). The glycan 3D structures are generated by the GLYCAM Web server developed by the Woods group (http://glycam.org)(*69, 70*). Table S1 and Fig. S4 show the glycans.

Missing hydrogen atoms were added to all systems, after which they were solvated in a (220 Å)^3^ and (215 Å)^3^ water box, with and without glycans, respectively. We added sodium (Na^+^) and Chloride (Cl^−^) ions to achieve a salt concentration of 150 mM. We used Visual Molecular Dynamics (VMD)(*71*) to construct all protein systems. In total, we built six distinct systems: WT Omicron S protein with and without glycans, laboratory-engineered Omicron S protein with and without glycans, and laboratory-engineered D614G S protein with and without glycans.

Simulations were conducted in NAMD(*72*) using the CHARMM36m protein force field(*73*), CHARMM36 glycan force field(*74*), and TIP3P water model(*75*). We used hydrogen mass repartitioning (HMR) along with a uniform 4-fs time step(*76, 77*). The simulations were run at a constant temperature (310 K) and constant isotropic pressure (1 atm), maintained by a Langevin thermostat and piston(*78*), respectively. Long-range electrostatics were calculated at every time step using the particle-mesh Ewald method(*79*). We set a short-range cutoff for Lennard-Jones interactions at 12 Å, with a switching function starting at 10 Å. We added extra bonds to hold the three α-helix structures (residues 1141 to 1162) together, imitating how they are held together by the virus membrane envelope, which was not modeled here. We started with restraining protein backbones while equilibrating side chains and glycans for 1 ns. After the initial equilibration process, each of the six protein systems was simulated for 1.4 µs with three replicates per system, resulting in an aggregated total of 25.2 µs of simulation data.

The accessible surface area (ASA) was quantified using the "measure sasa" command in VMD(*71*). In line with a previous study(*23*), the ASA of the protein with and without glycans was measured separately. Subsequently, the glycan-shielded area was calculated by subtracting the ASA with glycans from the ASA without glycans. The ASA was evaluated every 30 ns throughout the simulation trajectories, and subsequently, the average values were computed.

Kernel density estimation (KDE) plots were constructed using the Seaborn library(*80*) in Python. For determining the position of the RBD with the respect to the spike, two collective variables are defined as follows: a distance is measured between the centers of mass for RBD-A (336–518) and SD1-B (531–592), and a dihedral angle is measured using the centers of mass from the domains RBD-A (336–518), SD1-A (531–592), SD2-A (593–677), and NTD-A (27–307).

## Acknowledgements

JCG acknowledges support from the CDC under contract 75D30123P17466. Computational resources were provided through the Extreme Science and Engineering Discovery Environment (ACCESS; TG-MCB130173), which is supported by the National Science Foundation (NSF; #2138259, #2138286, #2138307, #2137603, and #2138296). This work also used the Hive cluster, which is supported by the NSF (1828187) and is managed by the Partnership for an Advanced Computing Environment (PACE) at GT.

## Data availability

The datasets generated and analyzed in the scope of this study are available from the corresponding authors upon request.

## Notes

### Competing Interest Statement

The authors have declared no competing interest.

