## Supporting Information for "Enhanced surface accessibility of SARS-CoV-2 Omicron spike protein due to an altered glycosylation profile"

[1]National Center for Environmental Health, Division of Laboratory Sciences. Centers for Disease Control and Prevention (CDC), Atlanta, Georgia, USA.

[2]School of Physics, Georgia Institute of Technology, Atlanta, Georgia, USA.

[3]National Center for Immunization and Respiratory Diseases, Centers for Disease Control and Prevention (CDC), Atlanta, Georgia, USA.

†These authors made equal contribution to this work

*To whom correspondence should be addressed.

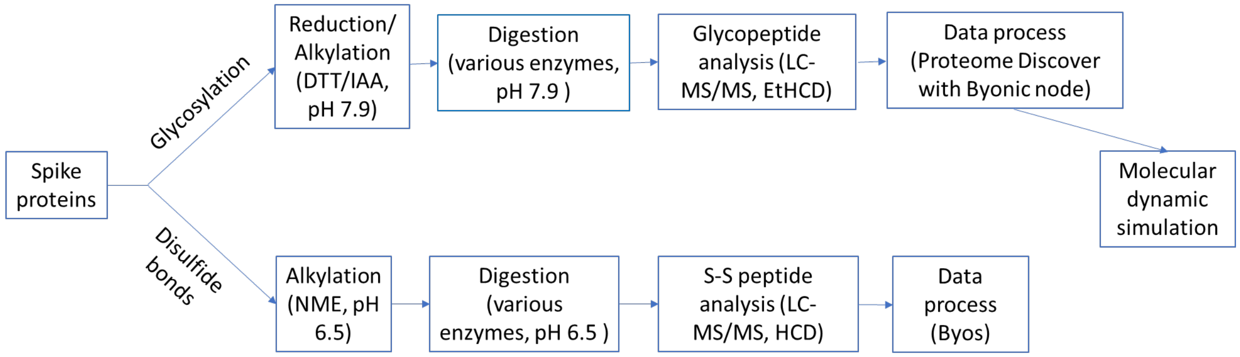


**Fig. S1. Workflow of the structural analysis of the SARS-CoV-2 Omicron and D614G spike proteins.**

Table S1. N-Glycosylation analysis data of SARS-CoV-2 Omicron and D614G S proteins obtained from the experiments of various digestion conditions. The top row of each sequon was selected for discussion in the text of this report.

| Sequon | S-Omicron | | | | | | S-D614G | | | | | | Digestion condition* |
|---|---|---|---|---|---|---|---|---|---|---|---|---|---|
| | Paucimannose | Oligomannose | Hybrid | Complex | Unoccupied | # Glycan | Paucimannose | Oligomannose | Hybrid | Complex | Unoccupied | # Glycan | |
| N17 | 14.6% | 2.0% | 3.0% | 80.4% | 0.0% | 39 | 4.9% | 0.7% | 2.8% | 91.6% | 0.0% | 34 | LysC-LysC |
| | 0.0% | 0.0% | 6.5% | 93.5% | 0.0% | 11 | 0.0% | 0.0% | 6.1% | 93.9% | 0.0% | 11 | LysC-aLP |
| | 0.0% | 0.0% | 4.6% | 95.4% | 0.0% | 10 | 0.0% | 0.0% | 4.5% | 95.5% | 0.0% | 10 | LysC-CHTR |
| N61 | 7.0% | 50.2% | 19.8% | 23.0% | 0.0% | 40 | 10.2% | 62.6% | 14.6% | 12.6% | 0.0% | 38 | LysC-aLP |
| | 1.6% | 51.3% | 18.6% | 28.5% | 0.0% | 32 | 2.0% | 68.9% | 14.0% | 15.1% | 0.0% | 28 | LysC-CHTR |
| | 2.8% | 54.3% | 18.8% | 24.1% | 0.0% | 35 | 4.5% | 67.4% | 14.3% | 13.8% | 0.0% | 33 | AspN-CHTR |
| N74 | 0.0% | 0.1% | 0.0% | 99.9% | 0.0% | 37 | 0.3% | 2.9% | 3.8% | 88.9% | 4.0% | 47 | LysC-CHTR |
| | 1.3% | 22.6% | 5.5% | 63.8% | 6.9% | 30 | 0.0% | 13.4% | 0.6% | 80.6% | 5.4% | 31 | AspN-CHTR |
| N122 | 1.0% | 61.4% | 30.5% | 7.1% | 0.0% | 43 | 1.5% | 71.0% | 22.2% | 5.3% | 0.0% | 44 | LysC-LysC |
| | 0.8% | 62.6% | 30.6% | 6.0% | 0.0% | 38 | 1.2% | 71.0% | 22.9% | 4.9% | 0.0% | 35 | LysC-Trypsin |
| | 2.7% | 52.3% | 30.5% | 14.5% | 0.0% | 32 | 2.6% | 57.4% | 20.5% | 19.4% | 0.0% | 30 | LysC-CHTR |
| N149 | 0.0% | 37.4% | 6.0% | 56.6% | 0.0% | 17 | 0.0% | 7.1% | 11.7% | 77.3% | 3.9% | 27 | AspN-CHTR |
| | 0.0% | 20.3% | 3.6% | 65.8% | 10.2% | 16 | 0.0% | 100.0% | 0.0% | 0.0% | 0.0% | 1 | LysC-Trypsin |
| | 0.0% | 27.6% | 4.7% | 63.8% | 3.9% | 14 | 100.0% | 0.0% | 0.0% | 0.0% | 0.0% | 1 | LysC-LysC |
| | 0.0% | 66.1% | 1.5% | 30.2% | 2.2% | 24 | 0.0% | 0.0% | 0.0% | 100.0% | 0.0% | 1 | LysC-aLP |
| N165 | 0.1% | 48.5% | 46.9% | 4.5% | 0.0% | 34 | 0.0% | 35.0% | 41.9% | 23.1% | 0.0% | 43 | LysC-Trypsin |
| | 0.0% | 35.5% | 57.6% | 6.8% | 0.0% | 32 | 0.1% | 27.9% | 51.8% | 20.2% | 0.0% | 45 | LysC-LysC |
| | 1.5% | 92.5% | 4.4% | 1.6% | 0.0% | 19 | 2.1% | 73.7% | 10.0% | 14.3% | 0.0% | 29 | LysC-CHTR |
| N234 | 0.4% | 99.5% | 0.0% | 0.1% | 0.0% | 10 | 0.0% | 99.4% | 0.3% | 0.3% | 0.0% | 12 | LysC-CHTR |
| | 0.0% | 100.0% | 0.0% | 0.0% | 0.0% | 4 | 0.0% | 100.0% | 0.0% | 0.0% | 0.0% | 6 | LysC-aLP |
| | 0.5% | 92.0% | 7.4% | 0.0% | 0.0% | 12 | 0.3% | 89.0% | 10.5% | 0.1% | 0.0% | 14 | LysC-Trypsin |
| N282 | 0.6% | 37.7% | 27.8% | 33.5% | 0.4% | 51 | 0.4% | 27.0% | 33.6% | 38.6% | 0.3% | 52 | LysC-LysC |
| | 0.8% | 33.5% | 24.5% | 40.9% | 0.3% | 51 | 0.7% | 24.5% | 34.4% | 39.4% | 0.9% | 51 | LysC-Trypsin |
| | 0.0% | 37.4% | 27.2% | 35.0% | 0.4% | 47 | 0.0% | 27.6% | 33.3% | 38.7% | 0.5% | 46 | LysC-CHTR |
| | 0.0% | 3.3% | 34.4% | 62.3% | 0.0% | 21 | 0.0% | 0.0% | 12.9% | 87.1% | 0.0% | 19 | LysC-aLP |
| N331 | 0.9% | 21.1% | 20.5% | 57.5% | 0.0% | 40 | 0.7% | 9.7% | 23.0% | 66.6% | 0.0% | 40 | LysC-aLP |
| N343 | 1.7% | 27.5% | 21.8% | 49.0% | 0.0% | 52 | 4.1% | 28.1% | 23.1% | 44.6% | 0.0% | 52 | LysC-aLP |
| N603 | 1.2% | 98.4% | 0.3% | 0.0% | 0.0% | 13 | 1.4% | 95.9% | 2.7% | 0.0% | 0.0% | 20 | AspN-CHTR |
| | 0.8% | 94.8% | 4.3% | 0.1% | 0.0% | 9 | 0.9% | 94.7% | 3.3% | 1.0% | 0.0% | 13 | LysC-CHTR |
| N616 | 2.2% | 47.1% | 27.8% | 22.9% | 0.1% | 44 | 2.7% | 59.5% | 28.5% | 9.2% | 0.1% | 37 | AspN-CHTR |
| | 0.5% | 51.2% | 27.7% | 20.6% | 0.0% | 31 | 0.8% | 65.0% | 28.6% | 5.6% | 0.0% | 24 | LysC-CHTR |
| N657 | 4.0% | 23.2% | 5.6% | 46.4% | 20.8% | 30 | 1.4% | 5.4% | 3.7% | 87.1% | 2.4% | 36 | LysC-CHTR |
| | 3.8% | 17.8% | 6.8% | 39.4% | 32.3% | 32 | 0.6% | 4.8% | 5.1% | 82.5% | 7.0% | 26 | LysC-aLP |
| | 0.1% | 16.9% | 4.1% | 55.5% | 23.4% | 31 | 0.0% | 7.8% | 1.0% | 78.7% | 12.5% | 13 | LysC-Trypsin |
| N709 | 1.6% | 97.9% | 0.2% | 0.3% | 0.0% | 10 | 2.2% | 93.4% | 1.6% | 2.9% | 0.0% | 17 | LysC-aLP |
| N717 | 1.3% | 97.2% | 1.5% | 0.0% | 0.0% | 9 | 1.8% | 95.7% | 2.5% | 0.0% | 0.0% | 10 | LysC-aLP |
| N801 | 0.6% | 99.1% | 0.3% | 0.0% | 0.0% | 12 | 1.3% | 96.8% | 0.8% | 1.1% | 0.1% | 23 | LysC-LysC |
| | 1.5% | 98.3% | 0.2% | 0.1% | 0.0% | 12 | 3.3% | 94.9% | 1.4% | 0.3% | 0.1% | 19 | LysC-aLP |
| | 1.1% | 97.9% | 0.3% | 0.5% | 0.1% | 12 | 2.5% | 66.8% | 1.0% | 29.6% | 0.1% | 26 | LysC-Trypsin |
| N1074 | 0.8% | 87.9% | 8.6% | 2.0% | 0.8% | 42 | 0.7% | 73.0% | 17.6% | 6.9% | 1.8% | 49 | LysC-LysC |
| | 0.9% | 84.6% | 10.7% | 2.5% | 1.3% | 32 | 0.4% | 77.4% | 14.3% | 6.1% | 1.8% | 37 | LysC-Trypsin |
| | 0.5% | 88.9% | 9.6% | 0.4% | 0.5% | 21 | 1.1% | 79.0% | 14.2% | 4.4% | 1.3% | 31 | LysC-CHTR |
| | 5.5% | 82.4% | 9.0% | 3.1% | 0.0% | 12 | 6.6% | 66.5% | 16.4% | 10.5% | 0.0% | 14 | LysC-aLP |
| N1098 | 0.2% | 46.5% | 46.4% | 6.1% | 0.8% | 43 | 0.3% | 31.1% | 57.6% | 10.2% | 0.8% | 50 | LysC-Trypsin |
| | 0.4% | 41.0% | 48.4% | 8.6% | 1.6% | 40 | 0.5% | 24.2% | 57.7% | 16.0% | 1.7% | 42 | LysC-CHTR |
| | 0.0% | 43.5% | 54.2% | 2.3% | 0.0% | 19 | 0.0% | 27.6% | 63.1% | 9.3% | 0.0% | 23 | LysC-LysC |
| | 2.1% | 37.4% | 49.1% | 8.3% | 3.1% | 39 | 1.9% | 21.4% | 58.6% | 14.7% | 3.4% | 41 | LysC-aLP |
| N1134 | 3.9% | 18.5% | 32.2% | 44.9% | 0.5% | 72 | 3.3% | 12.1% | 40.9% | 42.2% | 1.4% | 68 | AspN-CHTR |
| | 2.2% | 2.4% | 47.7% | 47.8% | 0.0% | 18 | 2.9% | 1.5% | 51.6% | 44.1% | 0.0% | 17 | LysC-aLP |
| | 0.0% | 46.5% | 25.7% | 27.8% | 0.0% | 12 | 0.0% | 37.1% | 25.1% | 37.8% | 0.0% | 12 | LysC-Trypsin |
| | 0.0% | 35.3% | 49.1% | 15.5% | 0.0% | 14 | 0.0% | 27.0% | 49.0% | 24.0% | 0.0% | 8 | LysC-CHTR |
| | 0.0% | 28.4% | 51.0% | 20.6% | 0.0% | 7 | 0.0% | 23.1% | 57.3% | 19.7% | 0.0% | 7 | LysC-LysC |
| N1158 | 1.6% | 9.2% | 6.9% | 81.9% | 0.4% | 30 | 0.7% | 2.3% | 3.6% | 92.6% | 0.8% | 30 | LysC-aLP |
| | 0.5% | 8.5% | 9.6% | 81.4% | 0.0% | 27 | 0.0% | 1.2% | 0.1% | 98.7% | 0.0% | 31 | AspN-CHTR |
| N1173 | 0.8% | 7.6% | 3.0% | 47.6% | 41.0% | 46 | 0.2% | 1.0% | 0.9% | 40.6% | 57.3% | 44 | AspN-CHTR |
| | 0.4% | 5.2% | 5.6% | 88.9% | 0.0% | 32 | 0.2% | 1.1% | 2.2% | 96.5% | 0.0% | 32 | LysC-aLP |
| N1194 | 0.0% | 5.6% | 2.0% | 56.3% | 36.0% | 45 | 0.0% | 1.0% | 0.6% | 46.1% | 52.3% | 40 | LysC-LysC |
| | 0.5% | 6.2% | 3.2% | 57.3% | 32.8% | 45 | 0.0% | 1.0% | 0.8% | 39.8% | 58.4% | 37 | LysC-Trypsin |

* The proteins were sequentially digested by listed combination of two enzymes with the first digestion conducted at 52$^{o}$C for 60 min and the second one at 37$^{o}$C overnight. CHTR and aLP represent chymtrypsin and alpha-litic protease, respectively.

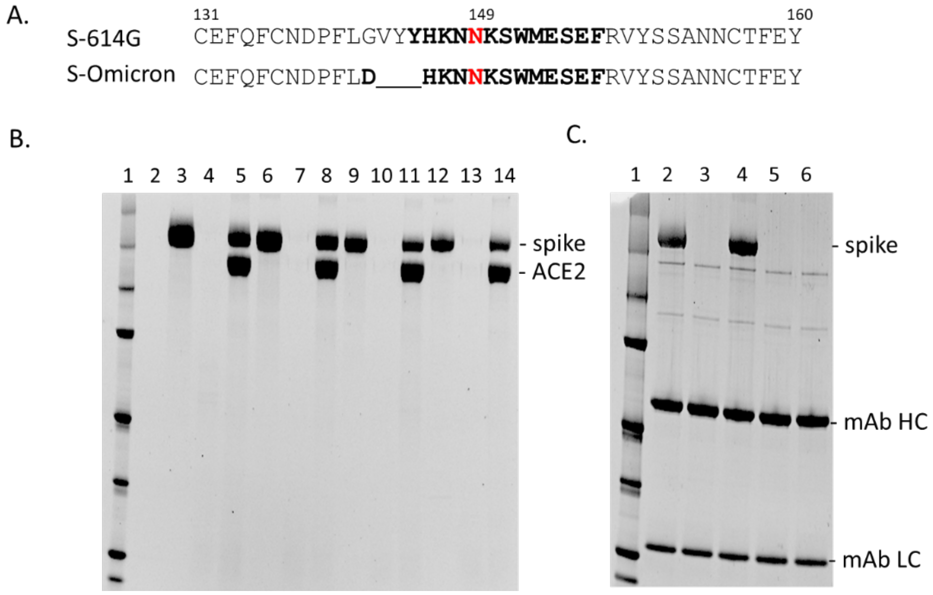


**Fig. S2.** (**A**) The region of the S-D614G and S-Omicron sequences containing the N149 sequon. Detected glycopeptides from these two proteins are displayed in bold. (**B**) Gel analysis of angiotensin-converting enzyme 2 (ACE2) binding to the wild type and N149Q mutants of the two spikes. Lane 1: marker; 2: empty; 3: S-D614G control in solution; 4: S-D614G and protein G beads; 5: S-D614G and ACE2 immobilized on protein G beads; 6: S-Omicron; 7: S-Omicron and protein G beads; 8: S-Omicron and ACE2 on protein G beads; 9: S-D614G-N149Q; 10: S-D614G-N149Q and protein G; 11: S-D614G-N149Q and ACE2-protein G beads; 12: S-Omicron-N149Q; 13: S-Omicron-N149Q and protein G; 14: S-Omicron-N149Q and ACE2-protein G beads. (**C**) Analysis of 4A8 mAb binding to the spike proteins. Lane1: marker; 2: S-D614G and mAb-protein G beads; 3: S-Omicron and mAb-protein G beads; 4: S-D614G-N149Q and mAb-protein G beads; 5: S-Omicron-N149Q and mAb-protein G beads; 6: mAb-protein beads control.

Table S2. Peak areas of the disulfide bonded (DB) peptides detected in both SARS-CoV-2 spike proteins of D614G and Omicron variants digested by various proteases.

| Digestion protease(s)[a] | DB #[b] | Peptide 1[c] | | Peptide 2 | | D614G | | Omicron | | Ratio (Omicron/ D614G) |
|---|---|---|---|---|---|---|---|---|---|---|
| | | Sequence | C1 posit. | Sequence | C2 posit. | Avg | SD | Avg | SD | |
| E3 | DB3 | DAVDCA | 291 | CTLK | 301 | 1.4E+09 | 4.0E+08 | 1.7E+09 | 6.4E+08 | 123% |
| E4 | DB3 | DAVDCAL | 291 | CTLK | 301 | 1.7E+09 | 4.2E+08 | 1.7E+09 | 2.6E+08 | 98% |
| E4 | DB3 | DAVDCALDPLSETK | 291 | CTLK | 301 | 8.1E+08 | 7.2E+07 | 5.0E+08 | 1.4E+08 | 62% |
| E4 | DB3 | DCAL | 291 | CTLK | 301 | 5.8E+08 | 2.5E+08 | 9.1E+08 | 1.7E+08 | 157% |
| E3 | DB3 | DCALDPLSET | 291 | CTLK | 301 | 9.7E+07 | 1.8E+07 | 1.5E+08 | 3.6E+07 | 158% |
| E3 | DB3 | DCALDPLSETK | 291 | CTLK | 301 | 9.3E+08 | 7.3E+07 | 1.0E+09 | 1.5E+08 | 109% |
| E4 | DB3 | DCALDPLSETK | 291 | CTLK | 301 | 6.6E+08 | 1.7E+08 | 4.1E+08 | 4.4E+07 | 62% |
| E2 | DB3 | YNENGTITDAVDCALDPLSETKCTLK | 291 | | 301 | 9.4E+06 | 2.5E+05 | 1.0E+07 | 2.7E+06 | 109% |
| E1 | DB4 | CPF | 336 | ISNCVADY | 361 | 1.2E+08 | 1.5E+07 | 3.9E+07 | 1.4E+07 | 32% |
| E1 | DB4 | CPFGEVF | 336 | ISNCVADY | 361 | 6.9E+07 | 1.9E+07 | 1.6E+07 | 4.2E+06 | 23% |
| E3 | DB4 | NLCPFG(D)EV | 336 | NCVA | 361 | 2.3E+08 | 1.8E+07 | 3.0E+08 | 6.3E+07 | 131% |
| E1 | DB5 | CY | 379 | KLPDDFTGCVIAW | 432 | 4.9E+08 | 1.8E+08 | 2.8E+08 | 7.2E+07 | 56% |
| E3 | DB5 | CYGVSPT | 379 | GCVIA | 432 | 3.4E+09 | 3.6E+08 | 2.3E+09 | 4.4E+08 | 68% |
| E1 | DB5 | KCY | 379 | LPDDFTGCVIAW | 432 | 5.0E+08 | 1.9E+08 | 2.3E+08 | 7.7E+07 | 46% |
| E1 | DB5 | KCY | 379 | KLPDDFTGCVIAW | 432 | 1.8E+09 | 1.8E+08 | 4.5E+08 | 7.8E+07 | 25% |
| E1 | DB6 | CF | 391 | HAPATVCGPK | 525 | 2.7E+08 | 2.5E+07 | 1.0E+08 | 3.6E+07 | 37% |
| E3 | DB6 | KLNDLCFT | 391 | TVCGPK | 525 | 1.6E+09 | 2.8E+08 | 1.7E+09 | 6.4E+08 | 105% |
| E3 | DB6 | KLNDLCFT | 391 | TVCGPKK | 525 | 2.7E+09 | 4.6E+08 | 1.9E+09 | 1.1E+08 | 71% |
| E3 | DB6 | KLNDLCFT | 391 | CGPK | 525 | 5.8E+08 | 1.2E+08 | 4.7E+08 | 4.9E+07 | 81% |
| E3 | DB6 | KLNDLCFT | 391 | CGPKK | 525 | 3.7E+08 | 7.0E+07 | 2.3E+08 | 2.7E+07 | 62% |
| E1 | DB6 | LNDLCF | 391 | HAPATVCGPKK | 525 | 2.7E+09 | 2.8E+08 | 1.1E+09 | 2.6E+08 | 43% |
| E3 | DB6 | LNDLCFT | 391 | TVCGPK | 525 | 6.4E+08 | 8.8E+07 | 8.7E+08 | 1.2E+08 | 137% |
| E3 | DB6 | LNDLCFT | 391 | TVCGPKK | 525 | 1.1E+09 | 1.8E+08 | 1.3E+09 | 1.6E+08 | 122% |
| E3 | DB6 | LNDLCFT | 391 | CGPK | 525 | 2.7E+08 | 4.2E+07 | 3.5E+08 | 5.8E+07 | 130% |
| E3 | DB6 | LNDLCFT | 391 | CGPKK | 525 | 2.4E+08 | 1.8E+07 | 2.7E+08 | 4.9E+07 | 111% |
| E2 | DB6 | LNDLCFTNVYADSFVIR | 391 | VVVLSFELLHAPATVCGPK | 525 | 1.3E+08 | 1.4E+07 | 1.1E+08 | 3.9E+07 | 83% |
| E1 | DB6 | NDLCF | 391 | HAPATVCGPK | 525 | 9.8E+08 | 1.1E+08 | 3.7E+08 | 6.5E+07 | 38% |
| E1 | DB6 | NDLCF | 391 | HAPATVCGPKK | 525 | 1.0E+08 | 2.1E+07 | 5.0E+07 | 1.3E+07 | 49% |
| E3 | DB8 | CV | 538 | PCSFGGV | 590 | 5.3E+08 | 3.5E+07 | 3.7E+08 | 7.4E+07 | 71% |
| E3 | DB8 | CV | 538 | PCSFGGVS | 590 | 8.2E+08 | 1.1E+08 | 5.5E+08 | 1.1E+08 | 67% |
| E1 | DB8 | CVNF | 538 | DITPCSF | 590 | 2.3E+09 | 8.1E+08 | 3.7E+09 | 1.3E+09 | 162% |
| E1 | DB8 | CVNF | 538 | EILDITPCSF | 590 | 1.5E+09 | 7.5E+08 | 3.4E+09 | 6.2E+08 | 218% |
| E3 | DB8 | CVNFNFNGLT(K) | 538 | PCSFGGV | 590 | 1.4E+09 | 3.7E+08 | 1.1E+09 | 1.1E+08 | 82% |
| E1 | DB8 | NKCVNF | 538 | DITPCSF | 590 | 1.8E+09 | 3.9E+08 | 1.5E+09 | 2.3E+08 | 80% |
| E1 | DB8 | NKCVNF | 538 | EILDITPCSF | 590 | 1.3E+09 | 2.9E+08 | 1.0E+09 | 2.4E+08 | 79% |
| E3 | DB10 | NNSYECDIPIGAGICAS | 662 | | 671 | 1.9E+08 | 2.9E+07 | 8.6E+08 | 8.8E+07 | 439% |
| E3 | DB11 | TSVDCT | 738 | FCT | 760 | 1.9E+08 | 4.1E+07 | 1.4E+08 | 2.9E+07 | 75% |
| E1 | DB11 | TSVDCTMY | 738 | CTQL | 760 | 2.2E+09 | 1.6E+08 | 2.6E+09 | 4.8E+08 | 119% |
| E1 | DB11 | TSVDCTMY | 738 | GSFCTQL | 760 | 3.3E+09 | 5.1E+08 | 4.7E+09 | 1.4E+09 | 142% |
| E1 | DB12 | ICGDSTECSNL | 743 | | 749 | 4.8E+09 | 4.6E+08 | 6.6E+09 | 1.7E+09 | 138% |
| E1 | DB12 | ICGDSTECSNLL | 743 | | 749 | 3.8E+08 | 4.3E+07 | 3.2E+08 | 3.4E+07 | 84% |
| E3 | DB12 | MYICGDSTECS | 743 | | 749 | 7.5E+08 | 3.6E+07 | 7.9E+08 | 4.9E+07 | 105% |
| E4 | DB13 | DCLG | 840 | DLICAQK | 851 | 7.8E+09 | 1.8E+09 | 7.6E+09 | 5.4E+08 | 98% |
| E1 | DB13 | GDCLGDIAAR | 840 | DLICAQK | 851 | 4.6E+08 | 1.5E+08 | 1.7E+09 | 4.6E+08 | 365% |
| E1 | DB13 | GDCLGDIAAR | 840 | DLICAQKF | 851 | 1.3E+09 | 3.4E+08 | 9.0E+08 | 1.6E+08 | 68% |
| E4 | DB13 | QYGDCLG | 840 | DLICAQK | 851 | 4.9E+08 | 1.9E+08 | 5.6E+08 | 9.8E+07 | 114% |
| E3 | DB13 | QYGDCLGDIAA | 840 | RDLICAQK | 851 | 2.1E+08 | 1.6E+07 | 3.4E+08 | 1.0E+08 | 162% |
| E2 | DB13 | QYGDCLGDIAAR | 840 | DLICAQK | 851 | 1.0E+10 | 1.3E+09 | 1.4E+10 | 4.2E+09 | 140% |
| E1 | DB13 | QYGDCLGDIAAR | 840 | DLICAQKF | 851 | 4.1E+08 | 1.7E+08 | 5.1E+08 | 7.8E+07 | 124% |
| E3 | DB13 | QYGDCLGDIAARDLICAQK | 840 | | 851 | 1.4E+10 | 2.4E+09 | 1.5E+10 | 3.4E+09 | 108% |
| E3 | DB14 | MSECV | 1032 | RVDFCGK | 1043 | 8.6E+08 | 5.2E+07 | 5.1E+08 | 9.5E+07 | 59% |
| E1 | DB14 | MSECVL | 1032 | CGK | 1043 | 2.0E+08 | 1.8E+07 | 1.1E+08 | 2.0E+07 | 59% |
| E1 | DB14 | MSECVL | 1032 | RVDFCGK | 1043 | 1.6E+09 | 3.0E+08 | 3.2E+09 | 6.0E+08 | 207% |
| E1 | DB14 | MSECVLGQSK | 1032 | RVDFCGK | 1043 | 2.8E+08 | 6.2E+07 | 3.0E+08 | 7.1E+07 | 111% |
| E2 | DB14 | MSECVLGQSK | 1032 | RVDFCGK | 1043 | 5.8E+09 | 3.6E+08 | 6.8E+09 | 6.1E+08 | 116% |
| E4 | DB14 | MSECVLGQSK | 1032 | RVDFCGK | 1043 | 6.5E+09 | 1.7E+09 | 4.2E+09 | 2.6E+08 | 65% |
| E2 | DB14 | MSECVLGQSK | 1032 | VDFCGK | 1043 | 5.0E+08 | 1.3E+08 | 5.2E+08 | 7.7E+06 | 105% |
| E4 | DB14 | MSECVLGQSK | 1032 | VDFCGK | 1043 | 2.0E+09 | 3.1E+08 | 1.8E+09 | 8.4E+07 | 91% |
| E2 | DB14 | MSECVLGQSKR | 1032 | VDFCGK | 1043 | 4.8E+09 | 2.3E+08 | 5.6E+09 | 6.5E+08 | 117% |
| E4 | DB14 | MSECVLGQSKR | 1032 | VDFCGK | 1043 | 5.3E+09 | 1.2E+09 | 5.1E+09 | 2.1E+09 | 96% |
| E2 | DB14 | MSECVLGQSKRVDFCGK | 1032 | | 1043 | 2.3E+08 | 2.5E+07 | 2.6E+08 | 2.0E+07 | 115% |
| E3 | DB15 | ICHDGK | 1082 | GNCDVV | 1126 | 2.1E+09 | 4.6E+08 | 1.9E+09 | 2.4E+08 | 91% |
| E3 | DB15 | ICHDGK | 1082 | SGNCDVV | 1126 | 1.5E+09 | 3.9E+08 | 1.1E+09 | 2.8E+08 | 75% |
| E3 | DB15 | ICHDGKA | 1082 | GNCDVV | 1126 | 3.0E+08 | 5.7E+07 | 2.0E+08 | 2.8E+07 | 66% |
| E3 | DB15 | ICHDGKA | 1082 | SGNCDVV | 1126 | 1.3E+08 | 3.3E+07 | 1.3E+08 | 1.0E+07 | 100% |
| E4 | DB15 | NFTTAPAICH | 1082 | DNTFVSGNC | 1126 | 9.6E+07 | 1.1E+07 | 3.4E+07 | 3.8E+06 | 36% |

a. enzyme(s): E1, E2, E3, and E4 represent trysin + chymotrypsin, Lys-C + trypsin, Lys-C + alphalitic protease, and Asp-N + trypsin, respectively.

b. The peptides contains cystine residue but do not form a disulfide bond are included. For example, DB3-free C1 represent a peptide with NME modified free Cys at 391 position 291.

c. The sequence and the amino acid position of the peptides are derived from the D614G spike. Letters in parentheses represent substitutions of Omicron spike protein.

A

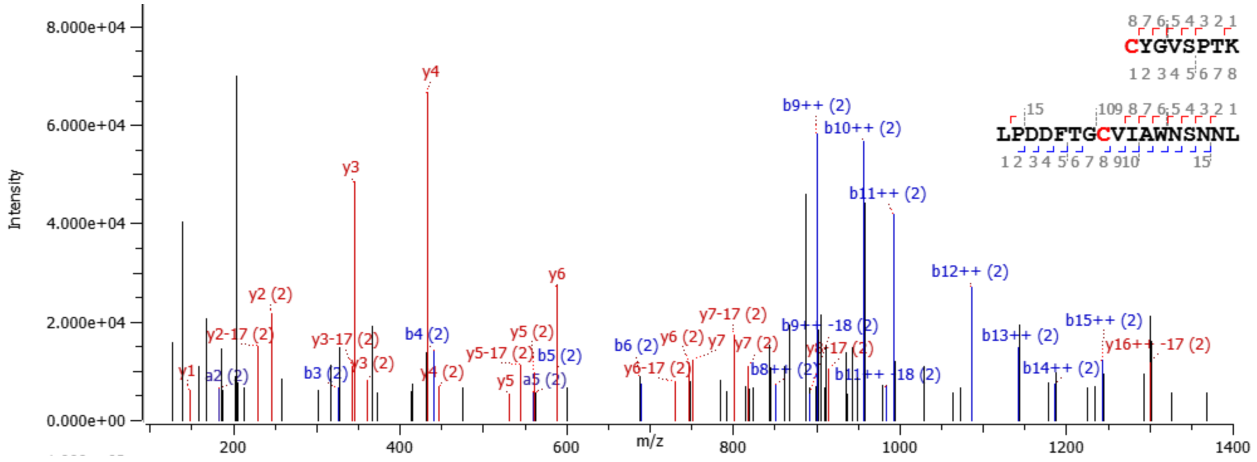


B

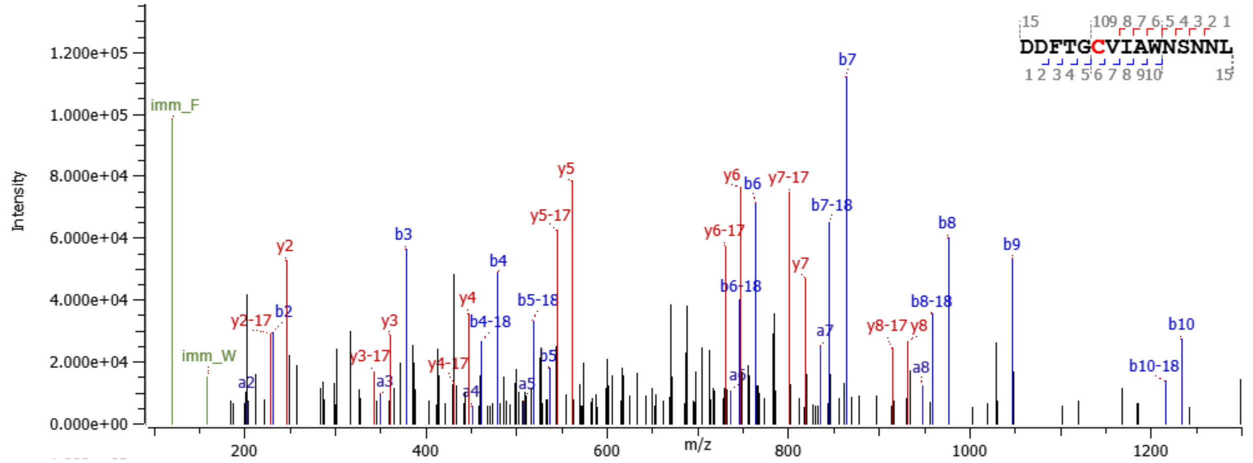


**Fig. S3.** MS/MS spectra of the C379-C432 peptide (**A**) and the peptide bearing the free C432 cysteine residue modified by NEM (**B**).

**Table S3. Glycan constituents at various positions in the 614G and Omicron S proteins.**

| Site | S-D614G | S-Omicron |
|---|---|---|
| N17 | N5H4A1 | N2H4F1 |
| N61 | N2H5 | N2H5 |
| N74 | N4H3F1 | N4H5F1A1 |
| N122 | N2H5 | N2H5 |
| N149 | N4H5F1A1 | N2H5 |
| N165 | N3H4 | N3H4 |
| N234 | N2H9 | N2H9 |
| N282 | N2H5 | N2H5 |
| N331 | N4H5F1A1 | N4H4F1 |

| | | |
|---|---|---|
| N343 | N2H5 | N2H5 |
| N603 | N2H5 | N2H5 |
| N616 | N2H5 | N2H5 |
| N657 | N4H4F1 | N2H5 |
| N709 | N2H7 | N2H8 |
| N717 | N2H7 | N2H7 |
| N801 | N2H7 | N2H8 |
| N1074 | N2H5 | N2H5 |
| N1098 | N3H5 | N2H5 |
| N1134 | N3H4F1 | N3H4F1 |
| N1158 | N5H3F1 | N5H3F1 |

* N, H, F, and A represent N-acetyl hexosamine (HexNac), hexose (Hex), fucose (Fuc), and N-acetylneuraminic acid (NeuAc) groups, respectively, in glycan compositions.

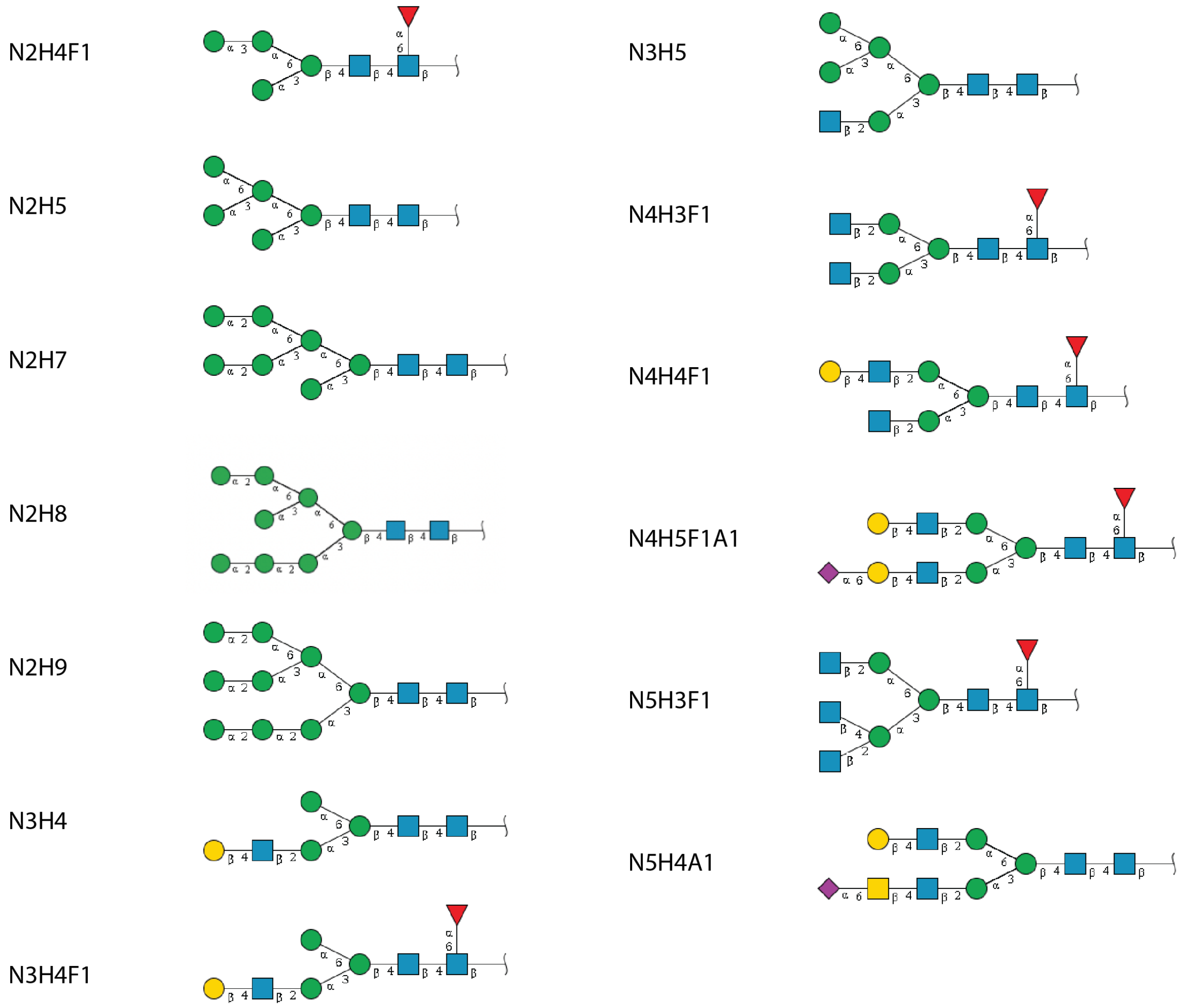

**Fig. S4. Specific glycan structures for various glycan constituents generated from the website GlyGen website (https://www.glygen.org)(*1*).**

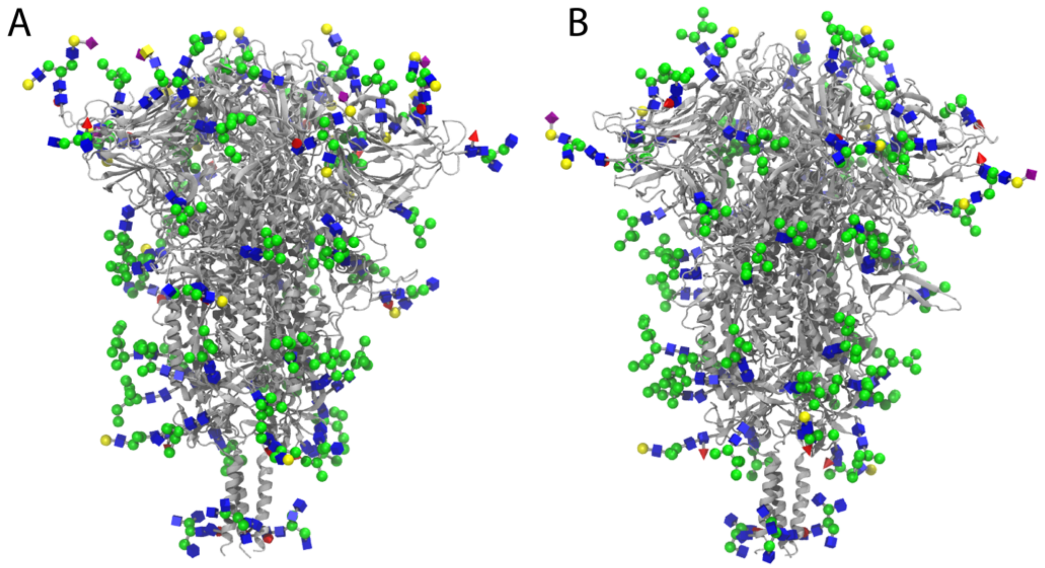


**Fig. S5. Constructed glycosylated SARS-CoV-2 spike proteins of D614G (A) and Omicron (B) variants.** The attached glycans are visualized using 3D-SNFG (symbol nomenclature for glycans) representations (*2*).

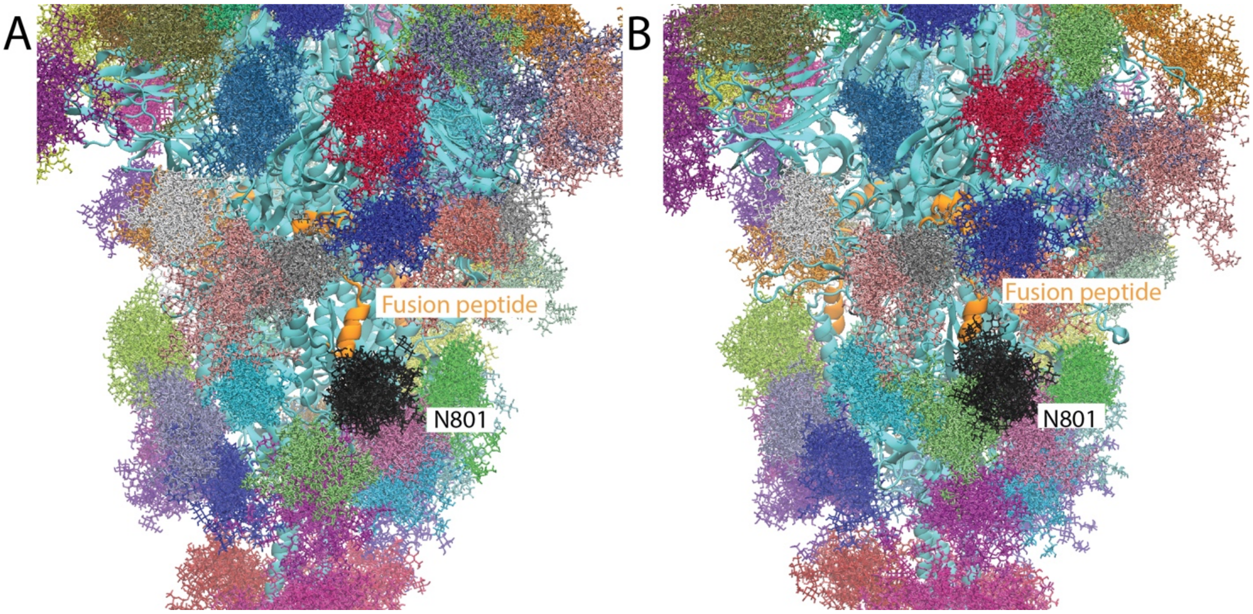


**Fig. S6. Glycan shielding near the fusion peptide.** The D614G (**A**) and Omicron (**B**) S proteins are depicted in cyan using a cartoon representation. The superimposed glycans are represented by colorful licorice models. These glycan configurations were captured at intervals of 0.25 μs throughout the net 4.2 μs of simulation trajectories for the D614G and Omicron S proteins.

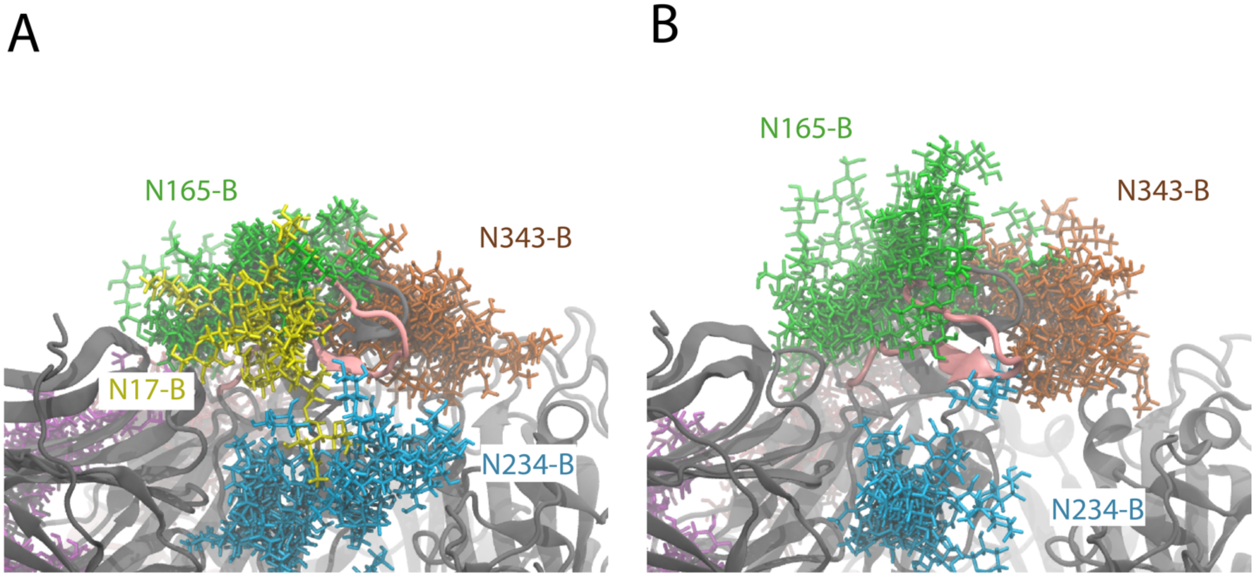


**Fig. S7. Glycan shielding of the receptor binding domain (RBD) in the SARS-CoV-2 S protein.** The residues from S469 to V483 of chain A in the D614G (**A**) and Omicron (**B**) S proteins are shown in pink. All glycan residues within 5 Å of the RBDs are depicted in the figures. These structures are superimposed at intervals of 0.25 μs along the respective simulation trajectories.

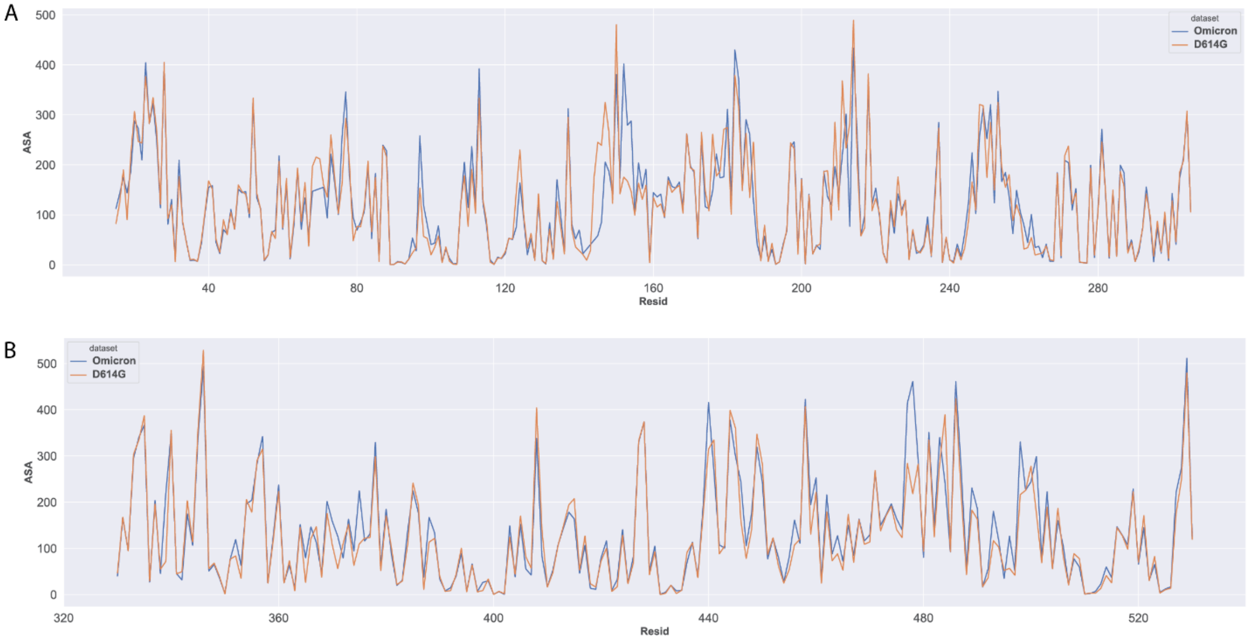

**Fig. S8. The accessible surface area (ASA) of the N-terminal domain (A) and receptor binding domain (B), factoring in the presence of glycans. The ASA is measured in Å$^2$.**

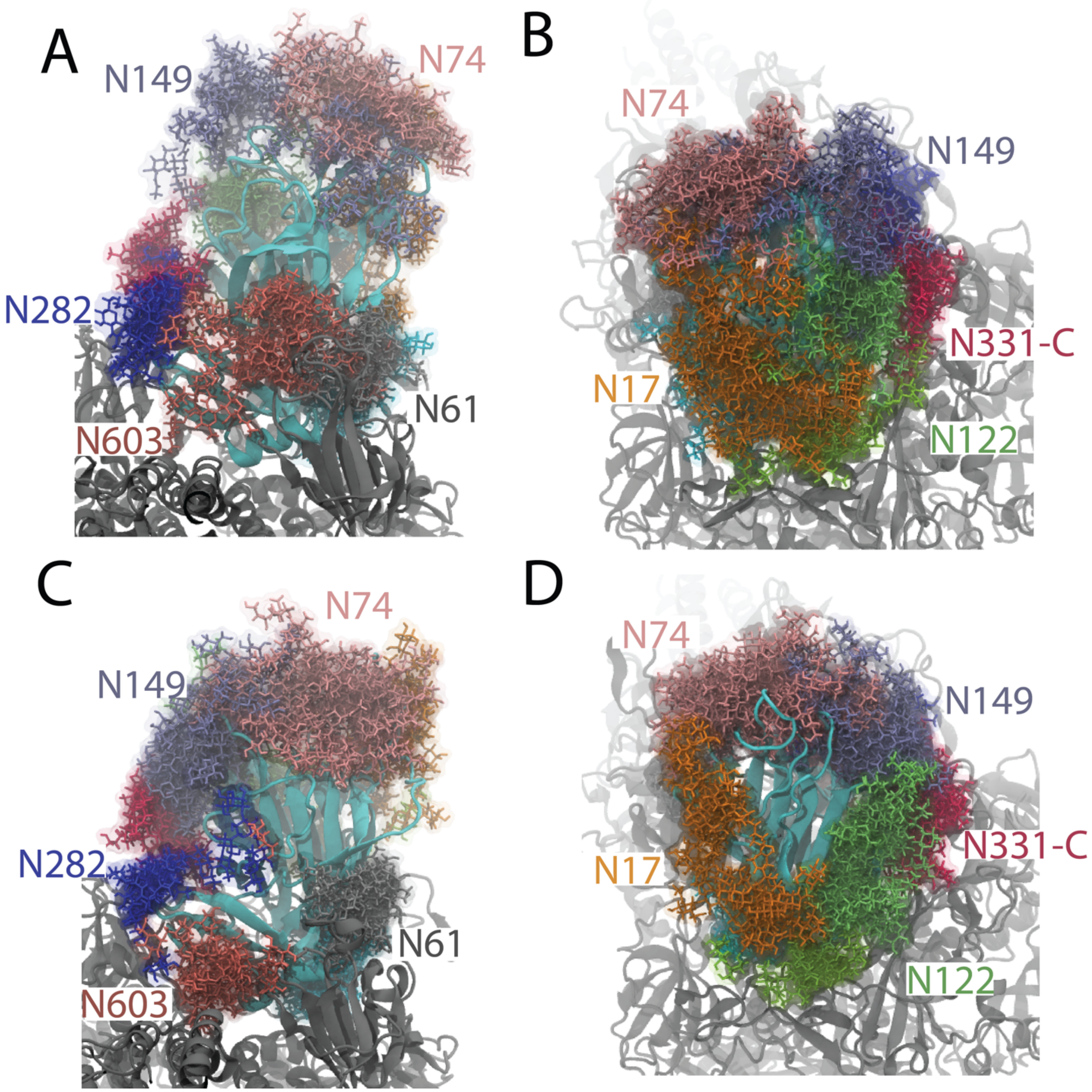


**Fig. S9. Glycan shielding of the N-terminal domain (NTD) in the SARS-CoV-2 S protein.** The NTDs of chain A in the D614G (**A, B**) and Omicron (**C, D**) S proteins are shown in cyan. All glycan residues within 5 Å of the NTDs are depicted in the figures. These structures are superimposed at intervals of 0.25 μs along the respective simulation trajectories.

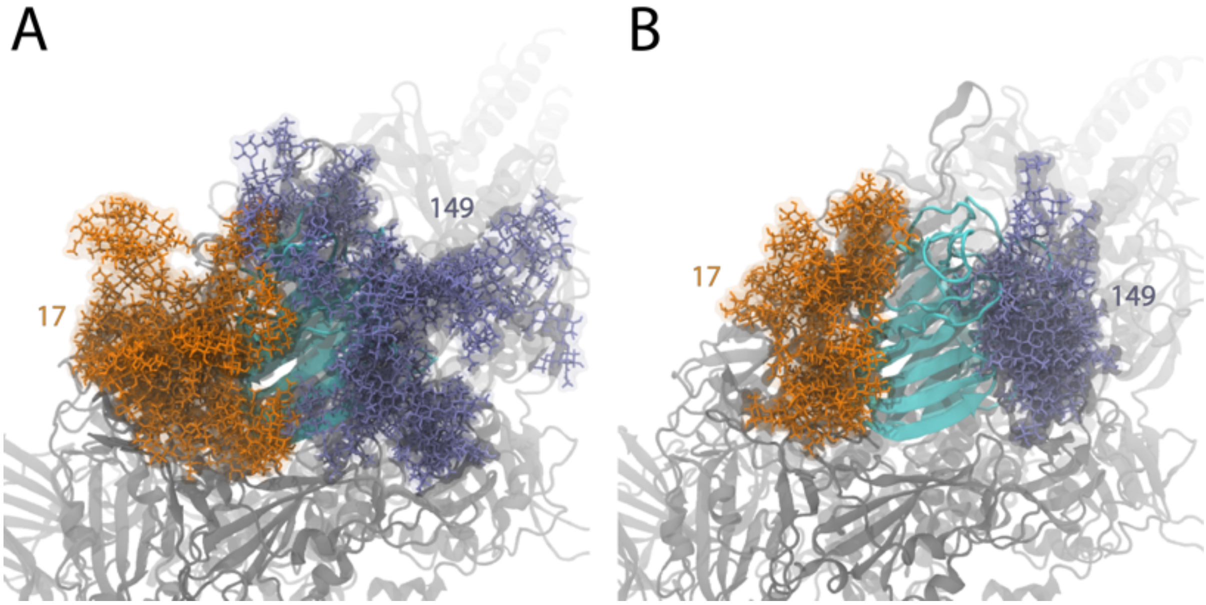


**Fig. S10. Full-length glycan coverage at positions N17 and N149.** Glycans are depicted in their complete length. The displayed structures are cumulative snapshots taken at 0.25-µs intervals along the respective simulation trajectories.
